# Global fitting for high-accuracy multi-channel single-molecule localization

**DOI:** 10.1101/2021.09.22.461230

**Authors:** Yiming Li, Wei Shi, Sheng Liu, Ulf Matti, Decheng Wu, Jonas Ries

**Affiliations:** Department of Biomedical Engineering, Southern University of Science and Technology, Shenzhen 518055, China; European Molecular Biology Laboratory, Cell Biology and Biophysics, Heidelberg 69117, Germany

## Abstract

Multi-channel detection in single-molecule localization microscopy (SMLM) greatly increases information content for various biological applications. Here, we present globLoc, a graphics processing unit (GPU) based global fitting algorithm with flexible PSF modeling and parameter sharing, to extract maximum information from multi-channel single molecule data. We show, both in simulations and experiments, that global fitting can substantially improve the 3D localization precision for biplane and 4Pi SMLM and color assignment for ratiometric multicolor imaging.

---

Single-molecule localization microscopy (SMLM) achieves nanometer superresolution and has become an important method for structural cell biology. Various extensions of SMLM using two or more detection channels are instrumental for this success, as they greatly increase information content that can be extracted from samples: Multi-color SMLM imaging of proteins labeled with fluorophores of different color can probe their spatial relations and interactions. It is usually realized using two spectral channels^1–3^ or one spatial channel combined with spectral detection in a second channel^4^. Three-dimensional (3D) SMLM techniques using two or more detection channels, such as biplane^5^ or multi-plane^6^ detection, self-bending point spread functions^7^ (PSFs), supercritical-angle fluorescence detection^8,9^ and multi-phase interference^10,11^, are powerful in investigating the intrinsic 3D organization of biological structures. Two or more fluorescence polarization channels are used to probe the orientation of fluorophores^12^, offering insight into the orientation of proteins in a molecular machinery. Recently, modulation enhanced localization microscopy that uses patterned excitation with rapid detection of different phases of the pattern on multiple parts of a camera, was used to increase the resolution of SMLM by a factor of two^13–16^.

Compared to the single-channel SMLM, data analysis for all these methods is complicated by the fact that measures from two or more channels have to be combined to result in the additional information (color, *z*-position, polarization state, interference phase, *etc*.). Typically, this is achieved by first fitting the fluorophores individually in each channel to extract corresponding parameters, and then combining the returned parameters from different channels to obtain the extra information^1-16^. Separate fitting of an individual fluorophore present in two channels is not optimal, as we neglect the information that the fitting parameters (*e*.*g*., 3D positions and photons) are highly correlated. If instead we were to use a global fitter that links the correlated parameters across different channels, this would decrease the number of fitting parameters, improve precision and robustness of the fit and avoid ambiguity when pairing corresponding parameters. Additionally, it would allow precise analysis of a fluorophore that is very dim in one of the channels and thus would escape molecule detection when fitted separately. In spite of the many benefits of analyzing separate channels simultaneously, global fitting is not widely used for the multi-channel single molecule localization. First approaches for global fitting^17–20^ lack flexibility with respect to the PSF models and fitting parameters. They are often designed for a specific imaging modality and difficult to be integrated into complete analysis workflows to be of general use.

Here, we developed globLoc, a general data analysis workflow and easy to use software for global fitting of single molecule data detected in separate channels. Its optimized analysis pipeline includes: generation of a precise transformation among the channels, calibration of a global multi-channel PSF, a GPU based global fitter that achieves maximum accuracy (**Supplementary Fig. 1, 2 and 3**) and ultra-fast fitting speed (**Supplementary Fig. 4**), as well as post-processing routines to extract the additional information (*z*, color, interference phase, polarization, *etc*.). Both, in simulations and on experimental data, we showed that global fitting indeed leads to a substantially improved localization precision for biplane and 4Pi-SMLM and color assignment in multi-color astigmatic SMLM.

We now give an overview of the globLoc analysis workflow (**Figure 1a, Supplementary Fig. 5**), and the details can be found in the **Methods**. We describe it using as an example dual-channel single molecule data. The extension to multi-channel data is straightforward. We first generate a global multi-channel experimental PSF model from image stacks of beads immobilized on a coverslip. To this end, we first calculate spline PSF models for each channel independently^21^ and fit each channel individually with the corresponding PSF model to obtain the precise bead positions. From corresponding bead positions in the two channels, we calculate the transformation between the channels. We then use cubic spline interpolation to register and average many bead stacks^21^, while keeping the fixed spatial relationship between the channels described by the transformation. Optionally, we re-calculate the transformation based on the actual SMLM experiment to account for channel drift. For this, we fit a sub-set of single molecule data in each channel separately using the corresponding PSF model and calculate the transformation based on the fitted coordinates. Besides using an experimental PSF model, our software also supports global fitting with a Gaussian PSF model.

**Figure 1:**
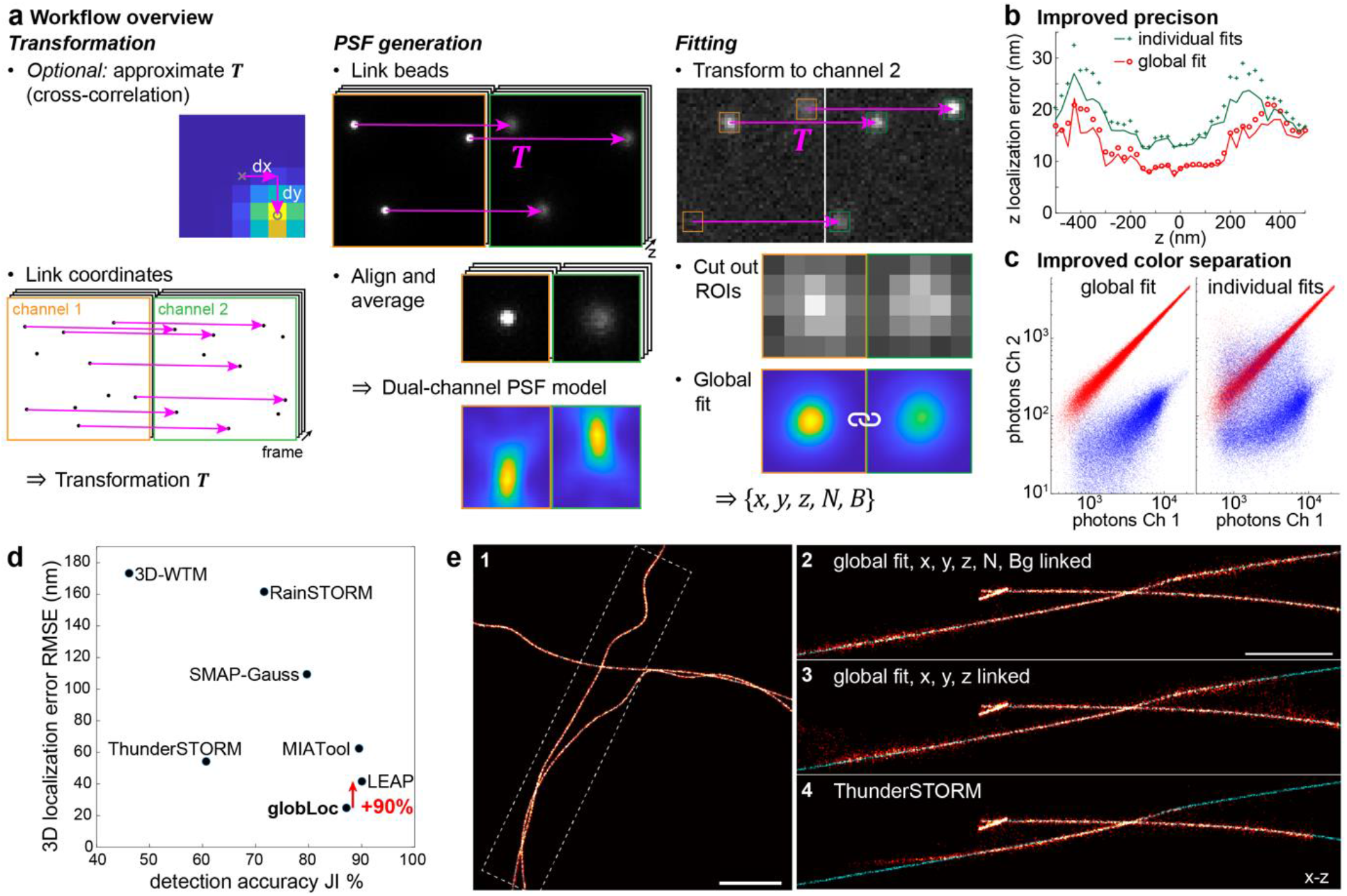
GlobLoc algorithm workflow and performance in simulated data. **a**, Overview of workflow of globLoc algorithm. The workflow includes 3 steps: calculation of multi-channel transformation using control points from beads or single molecule data; generation of multi-channel PSF models by properly averaging beads from multiple channels; fitting of the multi-channel single molecule data using global maximum likelihood estimation with flexible parameter linking. **b**, Comparison of the *z* localization error using globLoc and using single-channel fitting followed by CRLB-weighted averaging of positions. GlobLoc improved both the minimum theoretical localization uncertainty 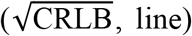 and the localization error (RMSE, circles) by approx. 1.5-fold compared to individual fitting (**Supplementary Fig. 1**). Simulations based on the biplane PSF model from the SMLM challenge 2016, 2500 detected photons in each channel and background of 20 photons per pixel. **c**, Comparison of color assignment for ratiometric multi-color imaging using globLoc and individual fitting. Photon numbers and background in each channel are based on experimental photon distributions (**Supplementary Fig. 7**) of AF647 (red) and CF680 (blue). **d**, Performance of different software on biplane data in the SMLM challenge 2016. GlobLoc (SMAP-global in the submission) outperformed all other software and improved the localization accuracy by 90% compared to the second-best algorithm LEAP^22^. **e**, Comparison of globLoc under different parameter sharing schemes and compared to ThunderSTORM (detailed settings in **Supplementary Fig. 10**) for biplane training data (MT0. N1. LD_BP) of the SMLM challenge 2016. **e2-4**, Side-view cross-sections of the region as indicated in e1 (e2: globLoc with *x, y, z*, photons and background photons shared; e3: globLoc with *x, y* and *z* shared; e4: ThunderSTORM.). Red-hot: fitted localizations; cyan: ground truth.

After calibration of the multi-channel PSF model and the transformation between different channels, the next step of the workflow is to perform global fitting to jointly analyze the multi-channel data using maximum likelihood estimation (MLE). On a standard GPU (NVIDIA RTX3090), our implementation reached ∼ 35,000 fits/s for regions of interest (ROI) with a size of 13×13 pixels, while the speed was ∼ 1,000 fits/s on a CPU (Intel Core i7-8700, **Supplementary Fig. 4)**. On simulated data for biplane SMLM, global fitting reached the Cramer-Rao-Lower-Bound (CRLB) in 3D over a large axial range (±600 nm, **Supplementary Fig. 1a**).

As globLoc is very flexible to link or unlink parameters between different channels, we compared the localization precision in the conditions of individual fit, global fit with only linking *xyz* positions and linking *xyz* positions plus photons per localization. Compared to individual fitting of the channels followed by CRLB-weighted averaging of positions^9^ (**Supplementary Note 1 and 2**), globLoc achieved about 1.5 times better *z* localization precision (**Figure 1b and Supplementary Fig. 1b**) and more robust parameter estimation (**Supplementary Fig. 1c**). This resolution improvement was further confirmed by participating in the continuously running 2016 SMLM Software Challenge^22^, in which globLoc improved the 3D localization precision by almost a factor of two on biplane data, compared to the second-best performing algorithm LEAP (**Figure 1d**). Our own comparison on the training data set (simulated microtubules) showed a clear improvement compared to the popular SMLM analysis software ThunderSTORM^23^ (**Figure 1e**). The improvement of globLoc compared to ThunderSTORM was even more apparent for when we analyzed experimental SMLM data of nuclear pore complex (NPC) protein Nup96, which we used as a reference standard^24^. In contrast to ThunderSTORM, globLoc was able to clearly resolve the two-ring structure of the NPC (**Figure 2a**). This is likely not only due to a better localization precision, but also an improved robustness of the fit.

**Figure 2:**
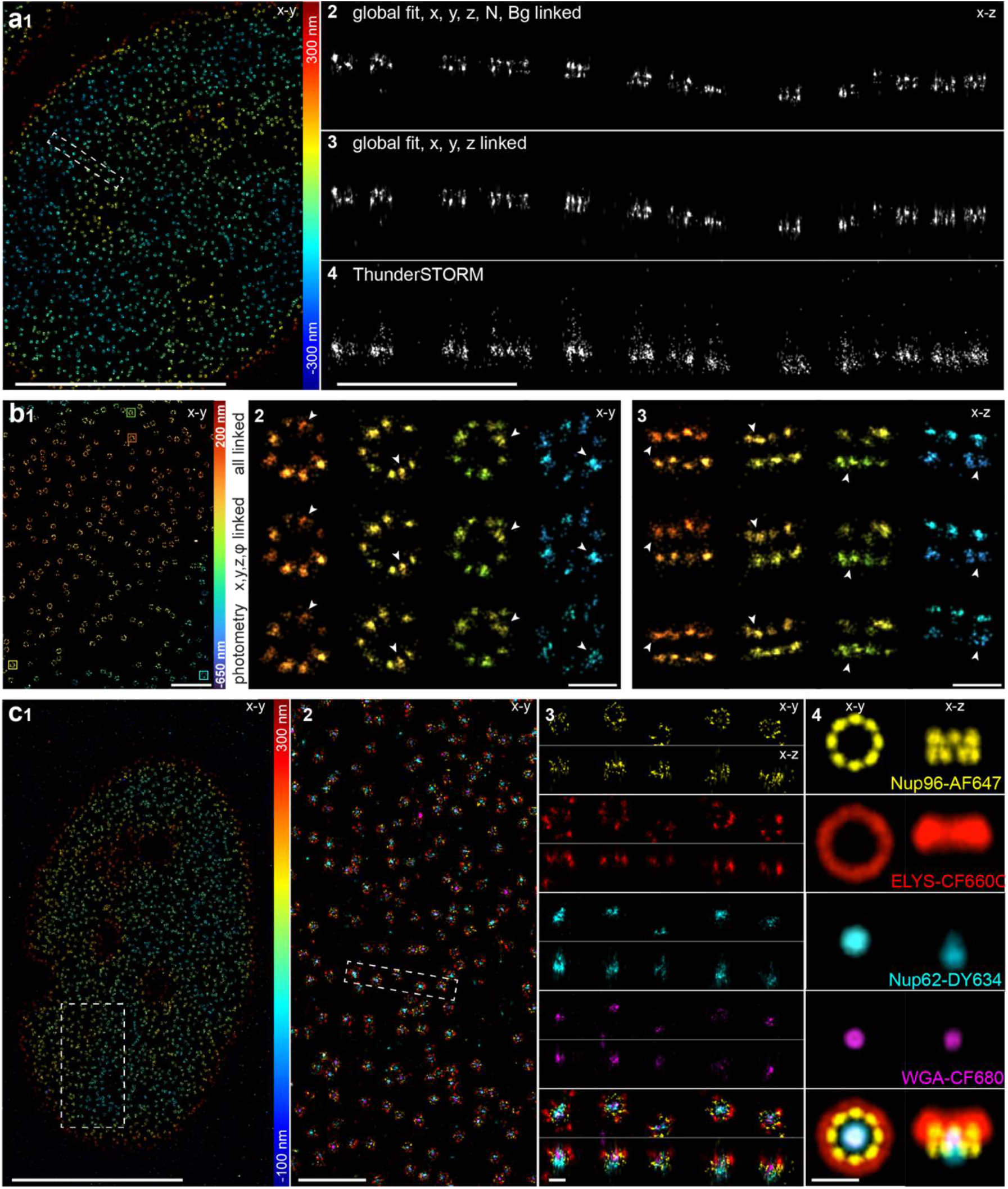
Demonstration of globLoc on biological structures. **a**, Comparison of globLoc on bi-plane data under different parameter sharing schemes and compared to ThunderSTORM (detailed settings in **Supplementary Fig. 10**), on the example of the nuclear pore protein Nup96-AF647. **a2-4**, Side-view cross-sections of the region as indicated in a1 (a2: globLoc with *x, y, z*, photons and background photons linked; a3: globLoc with *x, y* and *z* linked; a4: ThunderSTORM.). **b**, Comparison of globLoc on 4Pi-SMLM data under different parameter sharing schemes and compared to the state-of-the art analysis (photometry)^11^, on the example of Nup96-AF647. **b2-3**, Top-view and side-view reconstructions of selected NPCs at different *z* positions as indicated in b1 (all linked: globLoc with *x, y, z*, photons and background photons linked; *x,y,z, φ* linked: globLoc with *x, y* and *z* and the phase linked). **c**, 4 color 3D imaging of Nup62-DY634, Nup96-AF647, ELYS-CF660C and WGA-CF680 in the NPC using globLoc. **c1**, Top view of bottom nuclear membrane. **c2**, Zoomed image of the region indicated in d1. **c3**, Top view and side view images of the region indicated in c2. **c4**, Average of 200 NPC images by registering the Nup96 structures that we used as a reference. See **Supplementary Movie 1**. Scale bars 10 μm (a1, c1), 1 μm (a2-4, b1, c2), 100 nm (b2-3, c3-4).

GlobLoc is not limited to 2 channels. We implemented four-channel fitting for 4Pi-SMLM with multi-phase interference, using an advanced experimental 4Pi-PSF model that we developed recently^25^ (**Supplementary Note 3**). By fitting all four phase images globally with such a spline-interpolated experimental PSF model, globLoc achieved the CRLB in all dimensions, and greatly improved precision as well as accuracy compared to the state-of-the-art analysis (**Figure 2b** and **Supplementary Fig. 3**). As for biplane data, we found that additionally linking the photon number between different channels improved the localization precision in *z* by up to 1.5 times compared to only linking *xyz* (**Supplementary Fig. 3**).

Ratiometric multicolor SMLM images two or more dyes with overlapping emission spectra in two spectral channels and assigns the color of single molecules based on the relative number of photons detected in each channel (**Supplementary Fig. 6a and b)**. It has many advantages over conventional multicolor super-resolution imaging using dyes with well separated emission spectra^1,3,26^: 1) it has a negligible channel shift and chromatic aberration; 2) many of the best “blinking” dyes have similar emission spectra in the dark red range and are compatible with similar imaging conditions; 3) multi-color imaging can be performed simultaneously with one excitation laser. A key challenge for ratiometric color assignment is to precisely determine the photon number of the single molecules to distinguish their color. By using salvaged fluorescence reflected by the main excitation dichroic mirror, Zhang *et al*. have shown 3 color superresolution imaging of biological structures in 3D at 5-10 nm localization precision using 4Pi-SMS microscopy^3^. However, the salvaged fluorescence was only used for color assignment and did not contribute to the molecule localization.

Global fitting with globLoc improves the accuracy of determining the photons per localization (**Supplementary Fig 2b**) and thus the color assignment in both simulation and experiment (**Supplementary Fig. 6, 7 and 8**), while utilizing all detected photons for localization. To exploit our finding that linking photon numbers across channels increases the accuracy, we implemented a fitting approach in which globLoc fixes the relative photon numbers across the channels to different pre-calculated values and chooses the solution with the maximum likelihood (**Supplementary Fig. 7**). This approach reduced crosstalk during color assignment and minimized rejection of single molecules with close intensity ratios. It also makes a post-processing step for color assignment obsolete (**Supplementary Fig. 8**).

These innovations of GlobLoc enabled us to image and faithfully distinguish a record of 4 colors simultaneously in ratiometric 3D SMLM (**Figure 2c, Supplementary Fig. 9 and Supplementary Video 1**) and image Nup96, Nup62, Elys and WGA within single NPCs labeled with the dyes AF647, DY634, CF660C and CF680 with no apparent cross-talk. We averaged 200 NPC images by registering the Nup96 structures that we used as a reference^27^. This protein density map shows the average positions of the four NPC proteins, with Nup96 forming two rings with an 8-fold symmetry, Elys forming a large ring and Nup62 and WGA localizing at the central channel of the pore.

To summarize, we demonstrated that linking shared parameters during multi-channel single molecule localization substantially improves localization accuracy and reduces color assignment crosstalk. GlobLoc is fully integrated in SMAP^28^, allowing anyone to directly and easily use its full functionality (complete multi-channel calibration pipeline, versatile PSF model, flexible parameter sharing and fast fitting speed accelerated by GPU). In addition, as it is published as open-source with example code, it can also be easily used for custom software. We believe that globLoc will enable many groups to substantially improve their analysis workflows for multi-channel SMLM.

## METHODS

### Calculation of multi-channel transformation

Global fitting of multi-channel data relies on knowing the precise transformation among the channels. We developed a routine to calculate transformations from coordinates (bead positions or positions of single fluorophores) that we used during the generation of the multi-channel PSF model and for global fitting of multi-channel single molecule data. We describe our algorithm for a two-channel transformation (reference 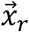 and target channel 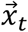). A multi-channel transformation is represented as several two-channel transformations from all target channels to the same reference channel. Our algorithm is as follows (**Figure 1a and Supplementary Fig. 5**):

1. Obtain approximate transformation ***T***_*0*_. This can be the transformation calculated in a previous experiment. Alternatively, we calculate it by first binning the coordinates in super pixels with a size of 50 nm. Then, we calculate the image cross-correlation and determine the position of the cross-correlation peak with sub-pixel accuracy from the position of the brightest pixel in a 4-fold upscaled image calculated by Gaussian filtering followed by cubic interpolation.
2. We transform all target coordinates to the reference channel using 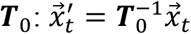.
3. We link coordinates in the reference and target channel if they are closer than a maximum distance 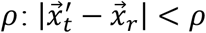. For fluorophores from SMLM experiments we only link coordinates from the same frame.
4. We calculate the precise transformation ***T*** based on the linked 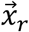 and 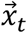 as anchor points. Usually, we use a projective transformation where ***T*** is represented by a 3 × 3 matrix, but we can use all transformations supported by Matlab, *e*.*g*., polynomial transformations.
5. If necessary, we repeat steps 3 and 4 with reduced *ρ*.

### Generation of multi-channel PSF models

Our algorithm to generate multi-channel PSF models from bead stacks is an extension of our work on generating single-channel experimental PSF models^21^. Again, we illustrate the steps of our algorithm on the dual-channel example, an extension to N channels is straight forward (**Figure 1a and Supplementary Fig. 5**):

1. We find candidate bead positions in each channel by calculating the mean image over all *z*-positions, Gaussian filtering and finding of local maxima above a user-defined threshold. These candidate positions are integers in the unit of camera pixels.
2. If no transformation among the channels exists, we first generate single-channel PSF models for each channel separately. We then fit the bead images using these new PSF models and finally use the fitted localizations to calculate ***T*** as described above.
3. We transform the coordinates of the candidate bead positions from the reference to the target channel: 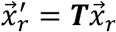. These target coordinates are continuous coordinates; thus we calculate the nearest integer pixel position by rounding the transformed coordinates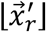 and calculating the shift between the rounded and original transformed coordinates 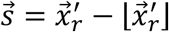.
4. We cut out ROIs around 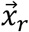 and 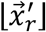 out of the bead stacks and shift the ROIs of the target channel by 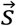 using cubic interpolation. If the target channel is mirrored with respect to the reference channel, we mirror the target ROI. This ensures that beads in both channels are shifted in the same direction during registration. Finally, we concatenate image stacks in both channels to form a single 3D array.
5. We create an initial template by averaging the 3D arrays over all beads and use 3D cross-correlation to register all beads to this template.
6. We reject those beads that have an insufficient overlap with the template (quality control) and calculate the next template as the average of the remaining shifted beads. We then register the central part of each bead to the new template.
7. We normalize the beads by the sum of the central slice of the reference stack.
8. We slightly filter the PSF models in *z* with a smoothing bspline and calculate a cspline representation for each channel.
9. To validate the PSF calibration, we fit each bead in the bead stack and compare the fitted *z* position with the true *z* position as denoted by the frame in the image stack.

### Extraction of multi-channel single molecule data

We implemented the workflow for global fitting of single molecule blinking events in the following way (again illustrated for two channels):

1. We calculate the global PSF model as described above from bead stacks.
2. Optionally, especially if we did not acquire bead stacks on the same day as the SMLM measurements, we calculate an improved transformation by fitting single molecule localizations in each channel independently and then using these localizations as anchor points to calculate ***T*** as described above. Otherwise, we use the transformation from the bead calibration.
3. We find candidate peaks in all channels using a difference of Gaussian filter and maximum finding. We then transform all candidates back to the first channel and average close-by candidate positions to obtain the coordinates of the candidates in the reference channel. Finally, we round to the nearest integer pixels to obtain 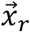.
4. As described for the beads, we transform the candidate positions to the target channels 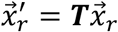 and calculate the shift between the rounded and original transformed coordinates 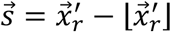.
5. Then we cut out ROIs around 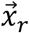 and 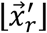. If the two channels are mirrored, we additionally mirror the ROIs and 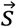.

### Maximum likelihood estimation of multi-channel single molecule data

We use a maximum likelihood estimator that jointly optimizes the combined likelihood across different channels. The objective function for MLE across different channels is given by:

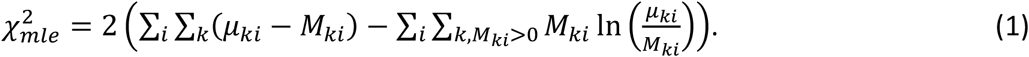

Here, *M*_*ki*_ is the measured photon number in the *k*th pixel of the *i*th channel. *μ*_*ki*_ is the expected photon number in the *k*th pixel of the *i*th channel. Similar to previous implementations^21,29,30^, we used a modified Levenberg-Marquardt (L-M) algorithm to minimize 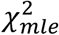 (**Supplementary Note 4**). For the multichannel nonlinear optimization process, the parameters can be classified as either shared (global) or non-shared (local) parameters, *θ* ∈ (***θ***_***p***,_ ***θ***_***qi***_). Here, ***θ***_***p***_ is the set of global parameters and ***θ***_***qi***_ is the set of local parameters of *i*th channel. The global parameters appear in all channels while the local parameters appear only in the individual channel. Depending on the imaging modality, any fitting parameter *θ* (*x, y, z/σ*_*PSF*_, photons, background) can be either linked as a global parameter among the channels or treated as a local parameter with different values in each channel. For global parameters, we define a transformation function to link parameters of different channels (translation and scale). The shared parameter in the *i*th channel can be written as: *θ*_*pi*_ *= S*_*pi*_*θ*_*p*_ *+ Δθ*_*pi*_. Here, *S*_*pi*_ and *Δθ*_*pi*_ are the scaling and translation factor, respectively. In this work, *Δθ*_*xi*_ and *Δθ*_*yi*_ are defined as the shifts 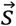 between the transformed ROI position and the actual ROI position which is rounded to integer pixels, as defined in item 4 of the section: Extraction of multi-channel single molecule data. The ratio of the photons between different channels *S*_*Ni*_, used for fixed photon ratio fitting, is determined from experimental single molecule data as the mean of the detected photons per localization for each dye. Therefore, the first derivative for a global parameter *θ*_*pi*_ in the *i*th channel is given by:

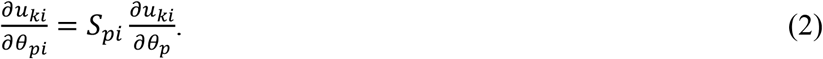

The first derivative for a local parameter *θ*_*qj*_ in the *i*th channel is 0 when *i* is not equal to *j*:

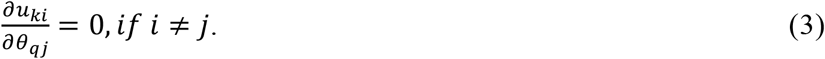

Therefore, the Jacobian matrix can be defined as:

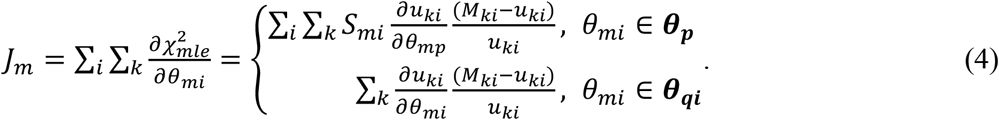

The Hessian matrix is defined as:

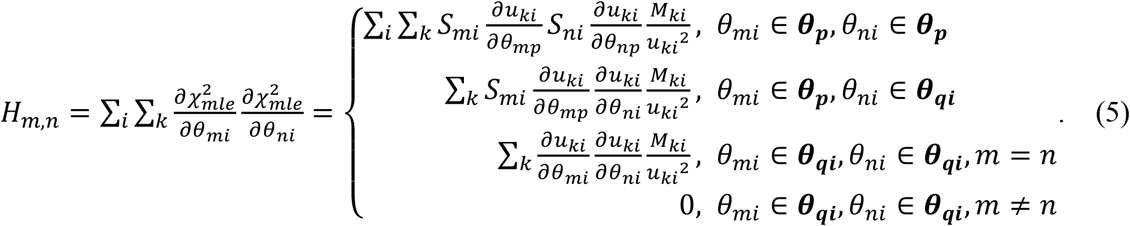

In the L-M algorithm, we updated the parameters by solving the linear equations: (***H*** *+ λ****I***)*Δ****θ*** *=* ***J***, with *λ* the damping factor and ***I*** a diagonal matrix equal to the diagonal elements of the Hessian matrix. The detailed algorithm can be found in **Supplementary Note 4**. Depending on different fitting modalities, we then calculate parameters of interest (*e*.*g*., color, polarization or z-position) from the fitted parameters (*e*.*g*., number of photons in each channel). Finally, we perform the usual post processing steps such as merging of localizations persisting over consecutive frames, drift correction and filtering based on log-likelihood and localization precision.

### Simulation and analysis of multi-channel data with experimental PSF models

For biplane data simulation (**Figure 1b and Supplementary Fig. 1**), the biplane experimental PSF model of the SMLM challenge 2016 was used. For each single molecule, we use 5,000 detected photons and 40 background photons for the simulation. The photons were then split into two channels with 1:1 ratio. 1,000 single molecule images were simulated for each *z* position (range from -600 nm to 600 nm) on a ROI of 15×15 pixels. Only Poisson noise was added to the images. The simulated biplane single molecule data was then fitted with three different schemes: 1) global fit with *x, y, z*, photons and background photons shared; 2) global fit with *x, y*, and *z* parameters shared; 3) individual fit for each channel and combination of the parameters of different channels with CRLB weighted arithmetic mean (**Supplementary Note 1**). The localization accuracy was calculated as the *root mean square error* (RMSE) of the fitted coordinates compared to the ground truth.

For the ratiometric astigmatic simulation (**Figure 1c, Supplementary Fig. 2, Supplementary Fig. 6** and **Supplementary Fig. 7**), a dual channel astigmatic experimental PSF acquired from multicolor beads was used. The photon distribution of these 4 different dyes were used for simulation: DY634, AF647, CF660C and CF680. The ratio of photons between two channels for the different dyes was determined from experimental data corresponding to **Figure 2c** as the mean of the detected photons per localization for each dye. We found photon ratios of *I*_2_/*I*_1_ *=* 0.39, 0.21, 0.07 and 0.02 for DY634, AF647, CF660C and CF680, respectively. Here, *I*_1_ and *I*_2_ are the photons from the bright and dark channels, respectively. For comparison of localization accuracy and CRLB (**Supplementary Fig. 2**), the photon ratio of 0.25 was used. 1,000 molecules with a ROI size of 15×15 pixels at each *z* position were used to calculate the RMSE. For ratiometric color separation (**Supplementary Fig. 6** and **Supplementary Fig. 7**), 50,000 single molecules were randomly placed at axial positions between -600 nm and 600 nm. The photon distribution of each dye follows the distribution of the experimentally acquired single molecules (**Supplementary Fig. 7a**). Three different methods were used to determine the color information: 1) The dual channel data was fitted separately; 2) The dual channel data was fitted globally with x, y and z as global parameters, photons and background as local parameters; 3) The dual channel data was fitted globally with x, y, z and photons as global parameters, background photons as shared parameter. The ratio of the photons between different channels was fixed during fit. For the first two methods, the color discrimination was realized by thresholding the normalized photon ratio: (*I*_1_ − *I*_2_)/(*I*_1_ *+ I*_2_). For the third method, the dual channel data was fitted with different fixed and pre-determined photon ratios of all 4 dyes between the two channels and we then chose the solution with the maximum likelihood.

For 4Pi single molecule data simulation (**Supplementary Fig. 3**), 2,000 photons/localization and 20 background photons/pixel were used for each objective. A full vectorial PSF model^31^ was used for simulations with the following parameters: NA 1.35, refractive index 1.40 (immersion medium and sample) and 1.518 (cover glass), emission wavelength 668 nm, astigmatism 100 mλ. 1,000 4Pi single molecule images with a ROI size of 15×15 pixels were simulated at each *z* position with four phase channels (0, π/2, π, 3π/2). The *x* and *y* positions are randomly distributed within -1 to 1 pixels around the center of each ROI. The simulated 4Pi single molecule data was then fitted with three different approaches: 1) global fit using IAB-based 4Pi-PSF model with *x, y, z*, phase, photons and background photons shared; 2) global fit using IAB-based 4Pi-PSF model with *x, y*, z and phase, parameters shared; 3) photometry based methods^11^.

### GPU implementation of globLoc and speed evaluation

We implemented the globLoc fitter with both spline and Gaussian PSF model using CUDA C/C++ in NVIDIA CUDA®-enabled graphic cards. The framework of the L-M iterative fitting method follows the previous work^21^. Each thread is pointed to a multi-channel single-molecule data and performs the entire fitting process for each single molecule. We put the single-molecule data in the global memory of the GPU and employed 64 threads for each block for the computation. Both the CPU and GPU based C++ code were compiled in Microsoft Visual Studio 2019 and called via Matlab 2019a (Mathworks) MEX files. For speed evaluation (**Supplementary Fig. 4**), we ran the CPU code on a personal computer using an Intel Core i7-8700 processor clocked at 3.2 GHz with 16GB memory. For the GPU-based evaluation, an NIVDA GeForce GTX 3090 graphics card with 24.0 GB memory was used.

### State-of-the-art workflows used for comparison

For biplane data analysis, we compared globLoc with the widely used ThunderSTORM software^23^. In the ThunderSTORM biplane analysis pipeline, a homography transformation is constructed from paired coordinates of the two channels. The biplane data is then fitted simultaneously using an astigmatic Gaussian PSF model. The detailed parameters used are shown in **Supplementary Fig. 10 and Supplementary Table 1**. For ratiometric multi-color assignment, we also compared to a workflow similar to that presented by Lehann et al (**Supplementary Fig. 9**)^26^. In short, localizations are fitted separately in the two channels. We then construct the transformation as described above and associate corresponding localizations in the two channels. Color assignment is then based on the relative fitted photon numbers.

### Cell culture

Before seeding of cells, high-precision 24 mm round glass coverslips (No. 1.5H, catalog no. 117640, Marienfeld) were cleaned by placing them overnight in a methanol:hydrochloric acid (50:50) mixture while stirring. After that, the coverslips were repeatedly rinsed with water until they reached a neutral pH. They were then placed overnight into a laminar flow cell culture hood to dry them before finally irradiating the coverslips by ultraviolet light for 30 min.

Cells were seeded on clean glass coverslips 2 days before fixation to reach a confluency of about 50 – 70% on the day of fixation. They were grown in growth medium (DMEM (catalog no. 11880-02, Gibco)) containing 1× MEM NEAA (catalog no. 11140-035, Gibco), 1× GlutaMAX (catalog no. 35050-038, Gibco) and 10% (v/v) fetal bovine serum (catalog no. 10270-106, Gibco) for approximately 2 days at 37 °C and 5% CO2. Before further processing, the growth medium was aspirated, and samples were rinsed with PBS (RT) to remove dead cells and debris. Unless otherwise stated, all experimental replicates were performed on cells of a different passage with separated sample preparation.

### Imaging Buffer

Glucose oxidase/catalase buffer supplemented with cysteamine (MEA) was used to image Nup96-SNAP-AF647-ELYS-CF660C-Nup62-DY634-WGA-CF680. GLOX+MEA contained 50 mM Tris/HCl pH8, 10 mM NaCl, 10% (w/v) D-glucose, 500 μg/ml glucose oxidase, 40 μg/ml glucose catalase and 35 mM MEA in H2O.

### Preparation of four-color NPC samples

Cells (Nup96-SNAP-tag, catalog no. 300444, CLS Cell Line Service, Eppelheim, Germany) on glass coverslips were prefixed in 2.4% (w/v) FA in PBS for 20 s before incubating them 10 min in 0.5% (v/v) Triton X-100 in PBS. Fixation was completed in 2.4% (w/v) FA in PBS for 20 min. FA was quenched for 5 min in 100 mM NH_4_Cl in PBS and then washed 3x 5 min in PBS. Fixed cells were blocked with Image-IT signal enhancer for 30 min and then incubated with 1 μM BG-AF647, 0.5% BSA and 1 mM DTT in PBS for 1 h to stain Nup96-SNAP-tag. Cells were washed 3x for 5 min with PBS and subsequently blocked with 5% (v/v) NGS (catalog no. PCN5000, lifeTech) in PBS for 1 h. Primary antibody labeling against ELYS was achieved by incubation with anti-AHCTF1 (HPA031658, Sigma-Aldrich) diluted 1:40 in 5% (v/v) NGS in PBS for 1 h. Coverslips were washed 3 times for 5 min with PBS to remove unbound antibody and subsequently stained with CF660C labeled anti-rabbit antibody (20183, Biotium, Fremont, CA) diluted 1:150 in PBS containing 5% (v/v) NGS for 1 h. After 3 washes with PBS for 5 min, the sample was postfixed for 30 min using 2.4% (w/v) FA in PBS, rinsed with PBS, quenched in 50 mM NH_4_Cl for 5 min and rinsed 3x 5 min with PBS. Labelling against Nup62 was performed by incubation with mouse anti-Nup-62 primary antibody (610498, BD Bioscience) diluted 1:50 in 5% NGS/PBS for 2h, 3x 5min washes of the coverslips with PBS and incubation over night at 4degC with 1:150 diluted secondary anti-mouse-DY634 antibody in 5%NGS/PBS. Unbound antibody was removed from the sample by washing 5 times with PBS. All incubations except otherwise stated were carried out at RT. Buffers used were also pre-equilibrated to RT.

Shortly before imaging, the sample was incubated for 10 min with 1:5000 diluted WGA-CF680 (29029-1, Biotium, Fremont, CA) in 100mM Tris, pH 8.0, 40mM NaCl, rinsed 3x with PBS and mounted onto a custom manufactured sample holder in imaging buffer. The holder was sealed with parafilm.

### Preparation of DY634-labelled secondary anti-mouse antibody

50ul of donkey anti-mouse IgG (H+L) (1,3mg/ml) (715-005-151, Dianova) was incubated with a 10-fold molar excess of DY634-NHS (634-01, Dyomics) in a final volume of 100ul PBS pH 7,4 overnight at RT. The labelled antibody was purified from free dye by running over an PBS equilibrated Zeba Spin desalting column (89889, Thermo Scientific) by gravity flow. Fractions containing the peak of the labelled antibody were identified by SDS-PAGE and pooled.

### Microscope setup

Single-objective SMLM image acquisition was performed at room temperature (RT, 24 °C) on a custom built microscope equipped with a high NA oil immersion objective (160x, 1.43-NA oil immersion, Leica, Wetzlar, Germany) described previously^32^. A commercial laser box (LightHub®, Omicron-Laserage Laserprodukte, Dudenhofen, Germany) equipped with Luxx 405, 488 and 638, Cobolt 561 lasers and an additional 640 nm booster laser (iBeam Smart, Toptica, Munich, Germany) were combined for wide field illumination. Lasers were focused onto a speckle reducer (LSR-3005-17S-VIS, Optotune, Dietikon, Switzerland) and coupled into a multi-mode fiber (M105L02S-A, Thorlabs, Newton, NJ, USA). The lasers were triggered using an FPGA (Mojo, Embedded Micro, Denver, CO, USA) allowing microsecond pulsing control of lasers. The output of the fiber was magnified by an achromatic lens and imaged into the sample plane. A laser clean-up filter (390/482/563/640 HC Quad, AHF, Tübingen, Germany) was placed in the excitation beam path to remove the fluorescence generated by the fiber. The focus of microscope was stabilized by a 785 nm infrared laser (iBeam Smart, Toptica, Munich, Germany) that was projected through the objective and reflected by the coverslip onto a quadrant photodiode, which was used as closed-loop feedback signal to the objective piezo stage (P-726 PIFOC, Physik Instrument, Karlsruhe, Germany). The fluorescence emission was filtered by a bandpass filter 676/37 (catalog no. FF01-676/37-25, Semrock) and then split into two channels (separated by ∼ 400 nm axially) using a 50:50 beamsplitter for biplane imaging with AF647. The astigmatic 3D imaging was acquired using a cylindrical lens (f = 1,000 mm; catalog no. LJ1516L1-A, Thorlabs) to introduce astigmatism. For astigmatic multicolor imaging with DY 634, AF647, CF660C and CF680, the fluorescence of the ratiometric multi-color imaging was split by a 665 nm long pass dichroic (catalog no. ET665lp, Chroma), filtered by a 685/70 (catalog no. ET685/70m, Chroma) bandpass filter for the transmitted light and a 676/37 (catalog no. FF01-676/37-25, Semrock) bandpass filter for the reflected light. An EMCCD camera (Evolve512D, Photometrics, Tucson, AZ, USA) was used to collect final fluorescence. Typically, we acquire 100,000 – 300,000 frames with 30 ms exposure time and laser power densities of ∼ 15 kW/cm^2^. The pulse length of the 405 nm laser is automatically adjusted to retain a constant number of localizations per frame.

4Pi-SMLM image acquisition was performed at RT based on an instrument as described previously^11^ with minor modifications. Two magnification matched silicone immersion objectives (1.35 NA, UPLSAPO, 100XS, Olympus) were used for better refractive index matching. The system was equipped with four excitation lasers: 405 nm (IBEAM-SMART-405 nm, 150 mW, Toptica), 488 nm (IBEAM-SMART-488-S-HP, 200 mW, Toptica), 560 nm (2RU-VFL-P-1500-560-B1R, MPB Communications, Pointe-Claire, Canada) and 642 nm (2RU-VFL-P-2000-642-B1R, MPB Communications). The excitation laser was filtered by a clean-up filter (390/482/563/640 HC Quad, AHF) and then reflected by a quadband dichroic (405/488/561/635, F73-867, AHF). The emission fluorescence was passed through the dichroic and then filtered by a quadband filter (432/515/595/730 HC, F67-432, AHF). The fluorescence was additionally filtered by a bandpass filter 676/37 (catalog no. FF01-676/37-25, Semrock) before collection on an sCMOS camera (ORCA-Flash 4.0v2, Hamamatsu). ∼ 200,000 images were acquired with 25 ms exposure time. The pulse length of the 405 nm laser is automatically adjusted to retain a constant number of localizations per frame.

## Supporting information

Tutorial of globFit

## Data availability statement

The experimental biplane and 4 color 3D astigmatic datasets can be freely downloaded from this website: https://www.embl.de/download/ries/globLoc/. All other data are available upon reasonable request from the corresponding authors.

## Code availability

Source code for the software used in this can be freely downloaded at https://github.com/jries/SMAP/tree/develop/fit3Dcspline/GlobLoc.

## Acknowledgements

We thank L. Zhou, S. Fu, J. Chen and M. Li for testing of the tutorials of globLoc. This work was supported by the Guangdong Natural Science Foundation Joint Fund (2020A1515110380 to Y. L.), Shenzhen Science and Technology Innovation Commission (Grant No. KQTD20200820113012029), the Startup grant from Southern University of Science and Technology, the European Research Council (grant no. ERC CoG-724489 to J.R.), the National Institutes of Health Common Fund 4D Nucleome Program (grant no. U01 EB021223 to J.R.), the Human Frontier Science Program (RGY0065/2017 to J.R.), the EMBL Interdisciplinary Postdoc Programme (EIPOD) under Marie Curie Actions COFUND (Y.L.), and the European Molecular Biology Laboratory.

## Author contributions

Y.L. and J.R. conceived the approach, developed the methods, wrote the software, and analyzed the data. W.S. and D.W. analyzed the biplane data and wrote the tutorial for globLoc. S.L. acquired and analyzed the 4Pi data. U.M. and Y.L. acquired the data. Y.L. and J.R. wrote the manuscript with input from all authors.

## Supplementary materials

**Supplementary Fig. 1.**
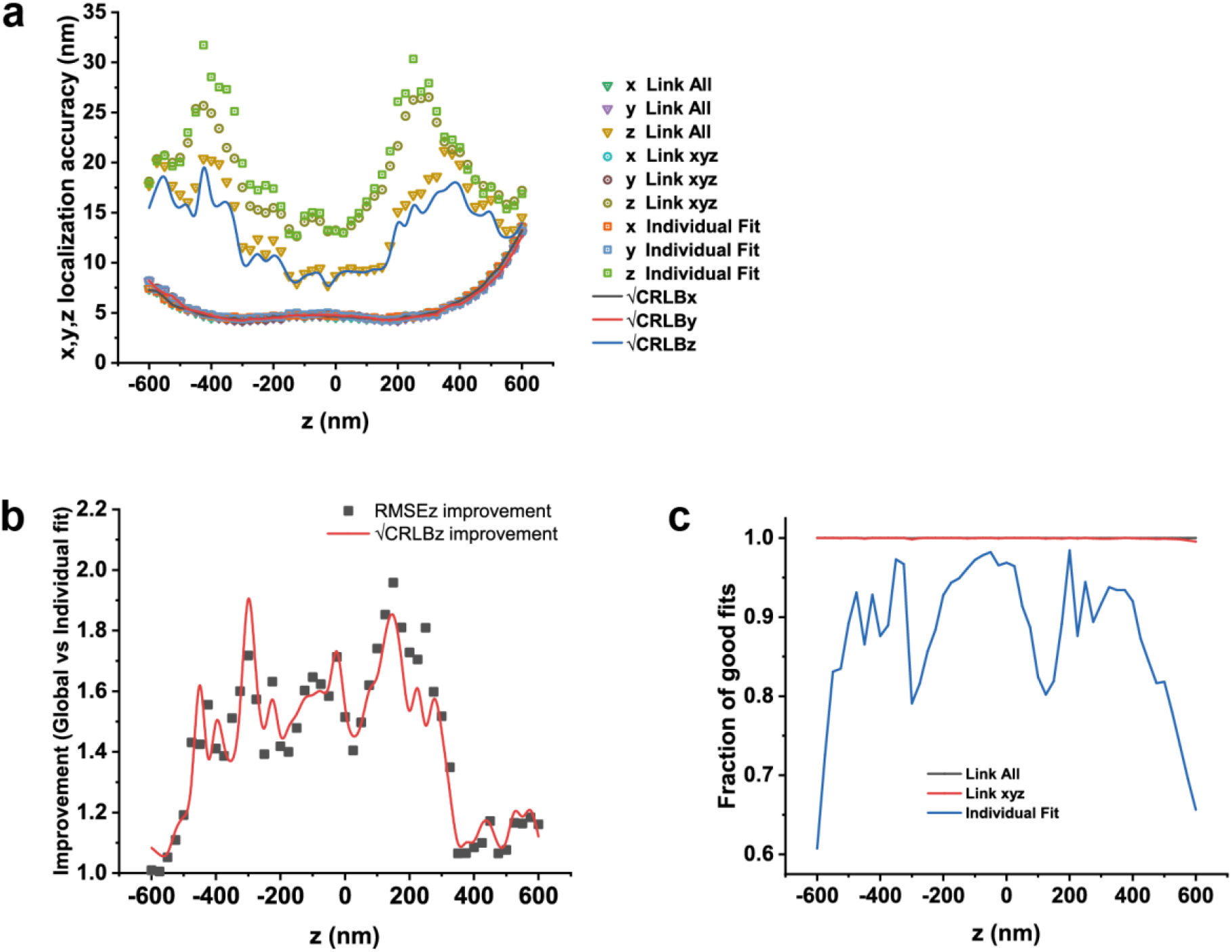
Performance of globLoc for biplane single molecule localization. Biplane single molecule images were simulated using an experimental PSF from the SMLM challenge 2016 with 2,500 photons/localization and 20 background photons/pixel in each channel. **a**, 3D localization accuracy (RMSE) of dual channel single molecule data using global fit and CRLB-weighted individual fit (Supplementary Note 1). By additionally linking the photons and background parameters during fitting, the *z* localization accuracy could be improved by more than 1.5 times compared to only linking *x, y*, and *z* positions. **b**, *z* localization accuracy improvement using global fit with all parameters linked compared to CRLB-weighted individual fit. **c**, The fraction of good fits for global fit and individual fit. If the distance between the returned *z* position and ground truth *z* position is within 8 times of the 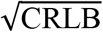 in the *z* direction, the fit is defined as a good fit. Only good fits were used to evaluate the localization accuracy.

**Supplementary Fig. 2.**
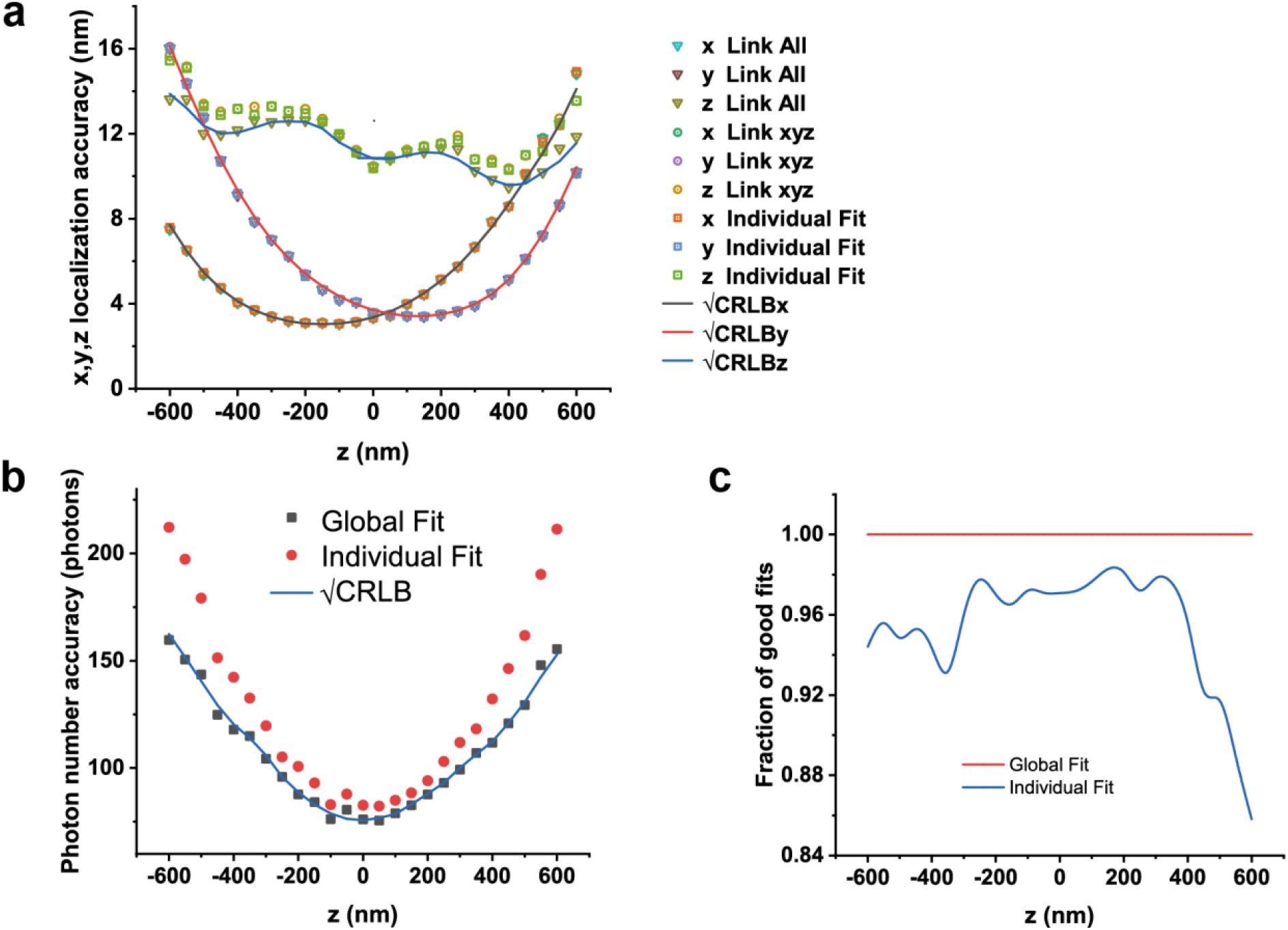
Performance of globLoc for dual color astigmatic single molecule localization. Dual channel single molecule images were simulated using an experimental dual channel astigmatic PSF with 4,000 photons/localization in one channel and 1,000 photons/localization in the other channel. 20 background photons/pixel were used for both channels. **a**, 3D localization accuracy (RMSE) of dual channel astigmatic single molecule data using global fit and CRLB-weighted individual fits (Supplementary Note 1). **b**, Photon number accuracy (RMSE) using global fit and CRLB-weighted individual fits. **c**, The fraction of good fits for global fit and individual fits. If the distance between the returned *z* position and ground truth *z* position is within 8 times of the 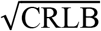 in the *z* direction, the fit defined as a good fit.

**Supplementary Fig. 3.**
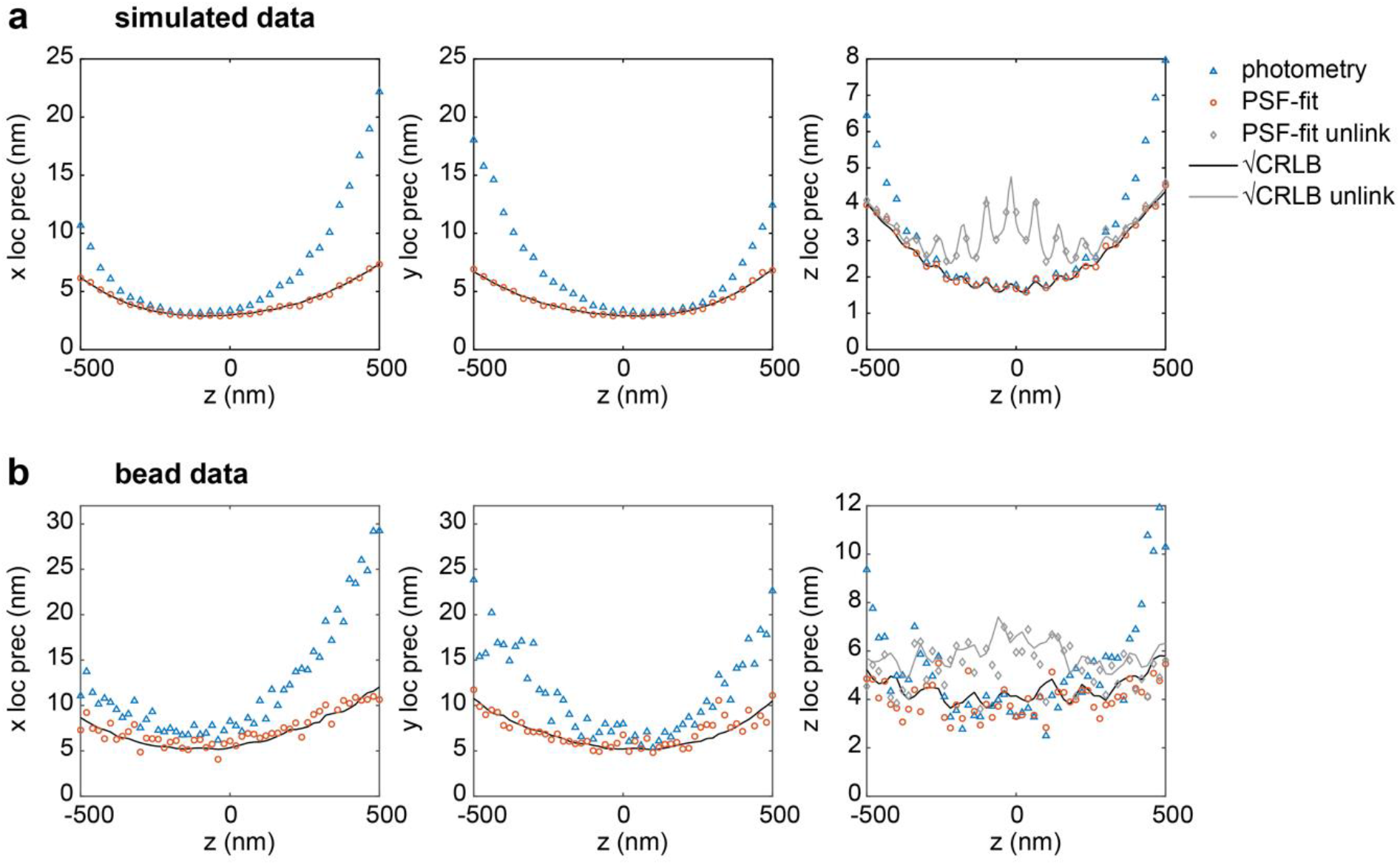
Performance of globLoc on 4Pi single molecule localization. Comparison of localization precisions and accuracies using an experimental 4Pi-PSF model fit and the photometry-based analysis method. **a**, Localization precision for simulated data. **b**, Localization precision for bead data. For simulated data, 1,000 4Pi single molecule images were simulated at each *z* position with four phase channels (0, π/2, π, 3π/2) with an astigmatism of 60 mλ and with *x* and *y* positions randomly distributed within -1 to 1 pixels around the center of each fitted region. For each objective, 2,000 photons/localization and 20 background photons/pixel were used. A full vectorial PSF model was used for simulations with parameters: NA 1.35. Refractive index 1.40 (immersion medium and sample) and 1.518 (cover glass). Emission wavelength 668 nm. For bead data, a single 40 nm red fluorescence bead (F8793, Invitrogen) was imaged at *z* positions from -500 nm to 500 nm with 20 nm step size and 37 frames were collected at each *z* position. The estimated photons/objective is 464 ± 40 (mean ± s.t.d) and the background photons/pixel is 0.47 ± 0.04 (mean ± s.t.d). The localization precision is calculated as the standard deviation. Theoretical estimation precision 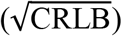 in *z* increases when unlinking the photon and background estimation in four phase channels.

**Supplementary Fig. 4.**
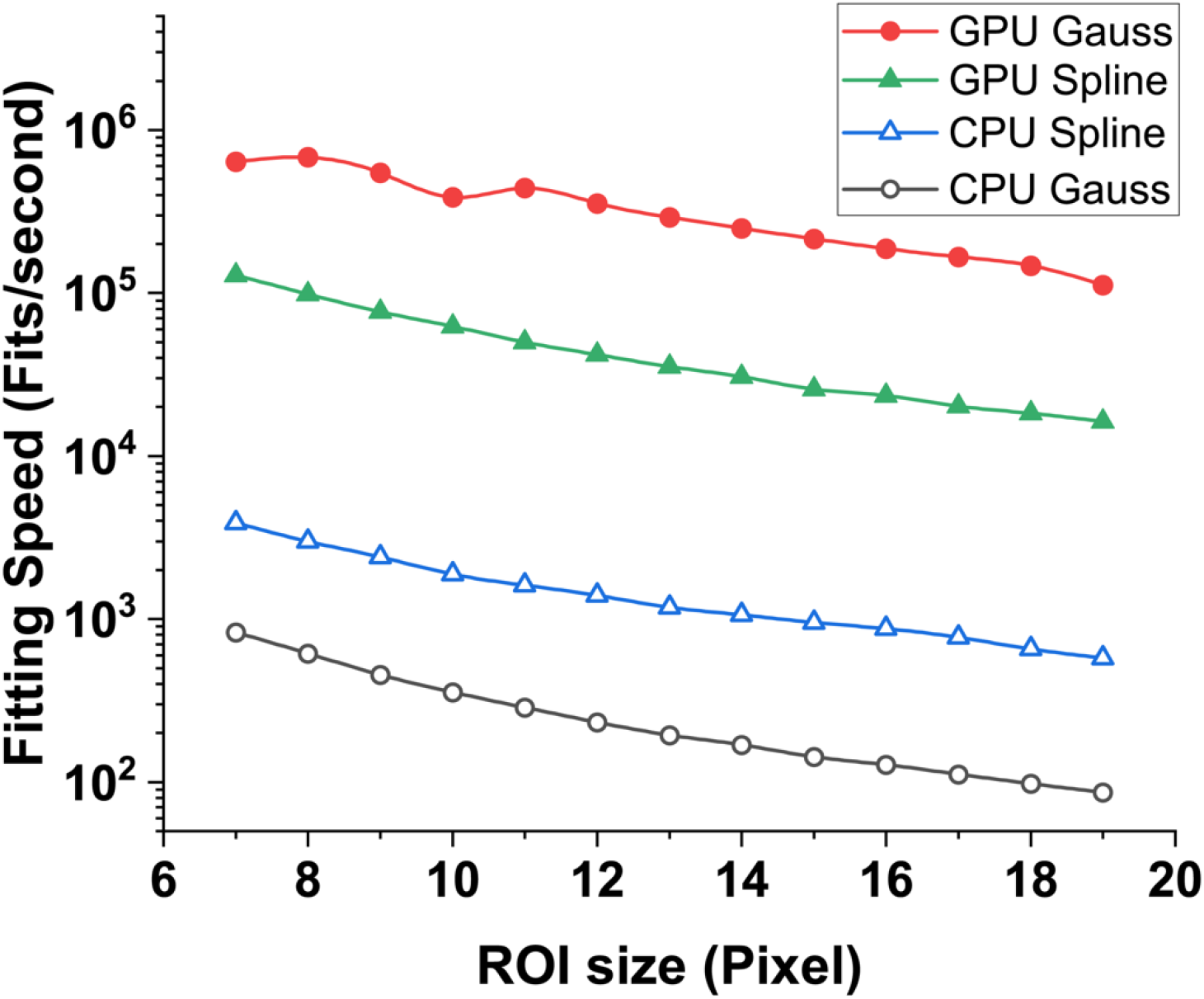
Computation speed of globLoc for Spline and Gaussian PSF model as a function of ROI size. Dual channel image data with the same ROI size in each channel was simulated for speed evaluation. Fits per second were measured on an i7-8700 CPU and a RTX3090 consumer graphics card. For fitting of the spline PSF model, The GPU code is overall about 30 times faster than the CPU code running on a single thread, while it is more than 1,000 times faster for fitting of Gaussian PSF model.

**Supplementary Fig. 5.**
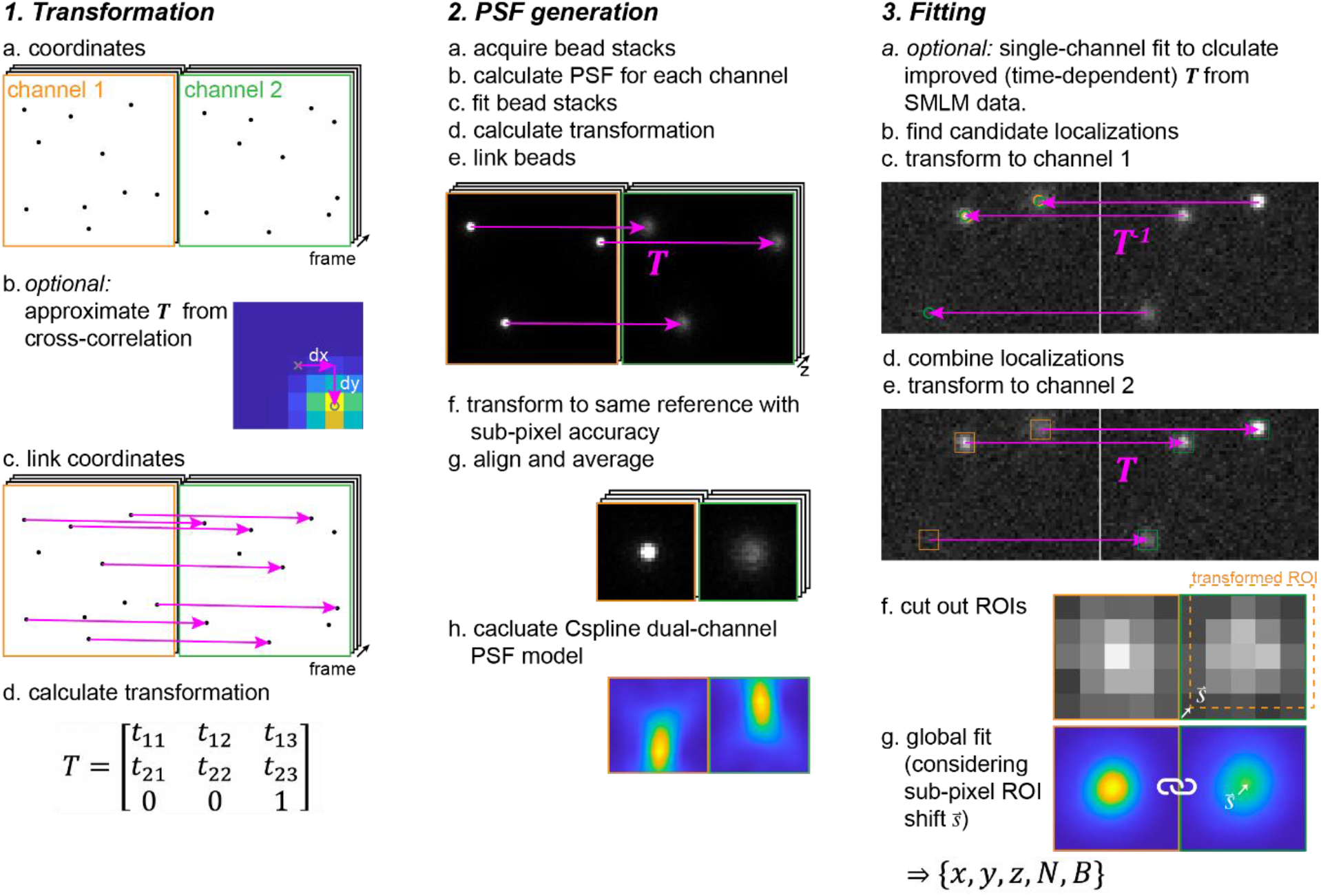
Extended workflow of globLoc. The overall workflow of globLoc can be divided into three parts: 1. Calculation of multi-channel transformation; 2. Generation of multi-channel PSF models; 3. global multi-channel maximum likelihood estimation. Detailed workflow can be found in **Methods**.

**Supplementary Fig. 6.**
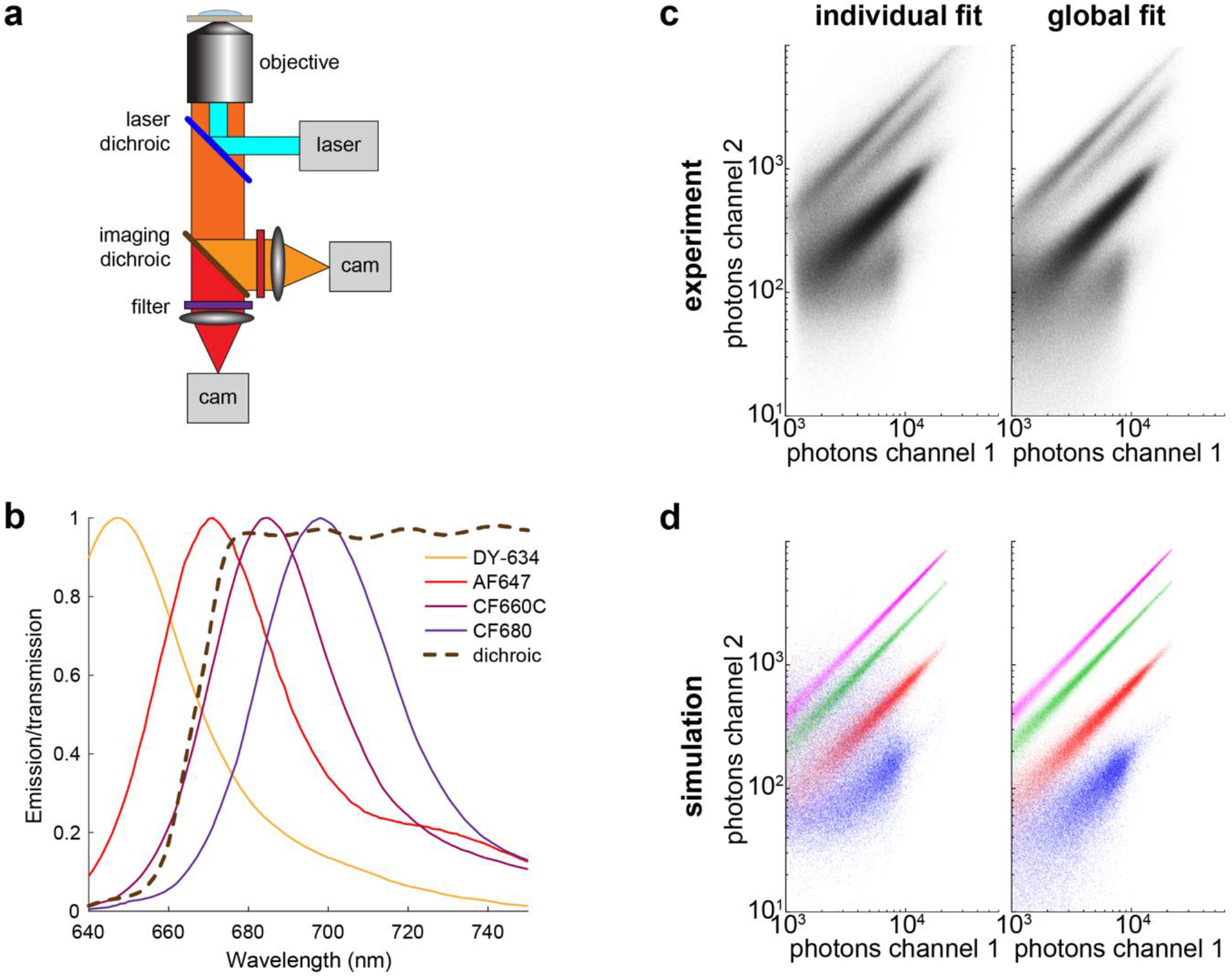
Photon distributions of global fit and individual fits. **a**, Simplified schematic of the ratiometric multicolor SMLM imaging system. The emission fluorescence is split into two channels by a dichroic mirror. The ratio of the single molecule fluorescence between these two channels are used to determine the color information of each single molecule. **b**, Emission spectra of DY-634, AF647, CF660C and CF680 and transmission profile of the dichroic beamsplitter. The experimentally determined ratio of photons between dark and bright channels for these four dyes are 0.39, 0.21, 0.07 and 0.02. **c**, Scatter plot of the fitted transmitted versus reflected photons per localization in a typical 4 color experiment (**Figure 2d**) using global fit and individual fit separately. **d**, Scatter plot of the fitted transmitted versus reflected photons per localization for simulated data using global fit and individual fit separately. Simulations were based on experimentally derived photon ratios and photon counts (**Supplementary Fig. 7a**).

**Supplementary Fig. 7.**
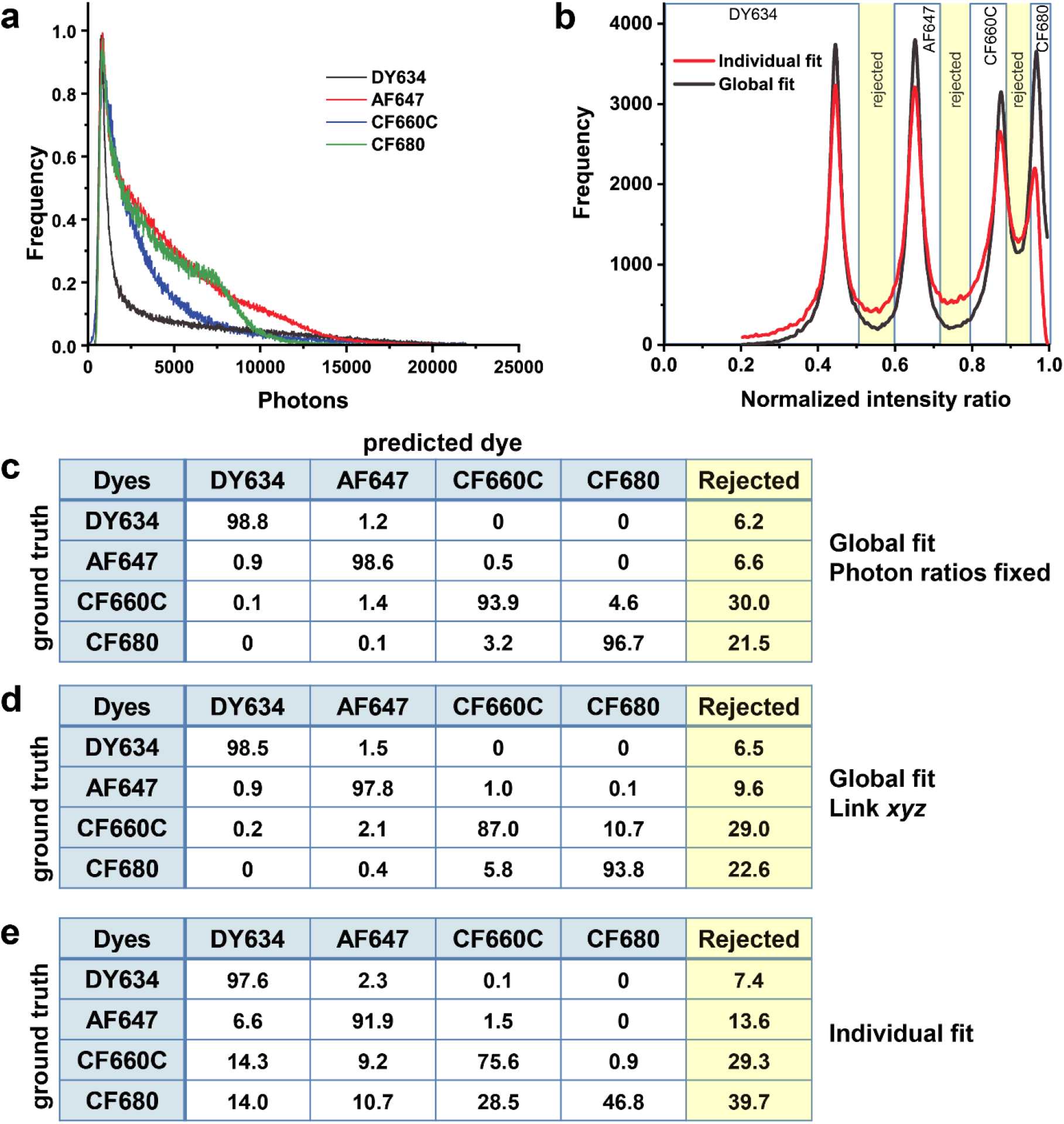
Cross-talk and rejected fraction for the 4 color ratiometric imaging using DY634, AF647, CF660C and CF680. **a**, Experimental photon distributions of DY634, AF647, CF660C and CF680 in our imaging system. The photon distribution of the simulated multicolor single molecules follows the experimental photon distribution for each dye. **b**, Comparison of normalized intensity ratio between two detection channels using global fit and individual fit. The normalized intensity ratio was calculated using the following equation: *r =* (*I*_1_ − *I*_2_)/(*I*_1_ *+ I*_2_). Here, *I*_1_ and *I*_2_ are the fitted photons per localization for the channel 1 and channel 2, respectively. The molecules were assigned to 4 different colors based on the intensity ratio threshold indicated by the 3 boxed regions (left to right: DY634, AF647, CF660C and CF680). **c**, The cross-talk (in %) of the 4 dyes using global fit with a fixed photon ratio between channels. The *x, y, z* and photon ratio were linked. The photon ratios were determined experimentally as indicated before. Each single molecule was fitted with 4 different photon ratios separately and the solution with the maximum likelihood was chosen. Here, we filtered out molecules for which the second highest likelihood was within 0.5% of the maximum likelihood. **d and e** are the cross-talk of the 4 dyes analyzed by a global fit, only linking *xyz*, and by fitting the two channels separately. The molecules were filtered and assigned as described in **b**.

**Supplementary Fig. 8.**
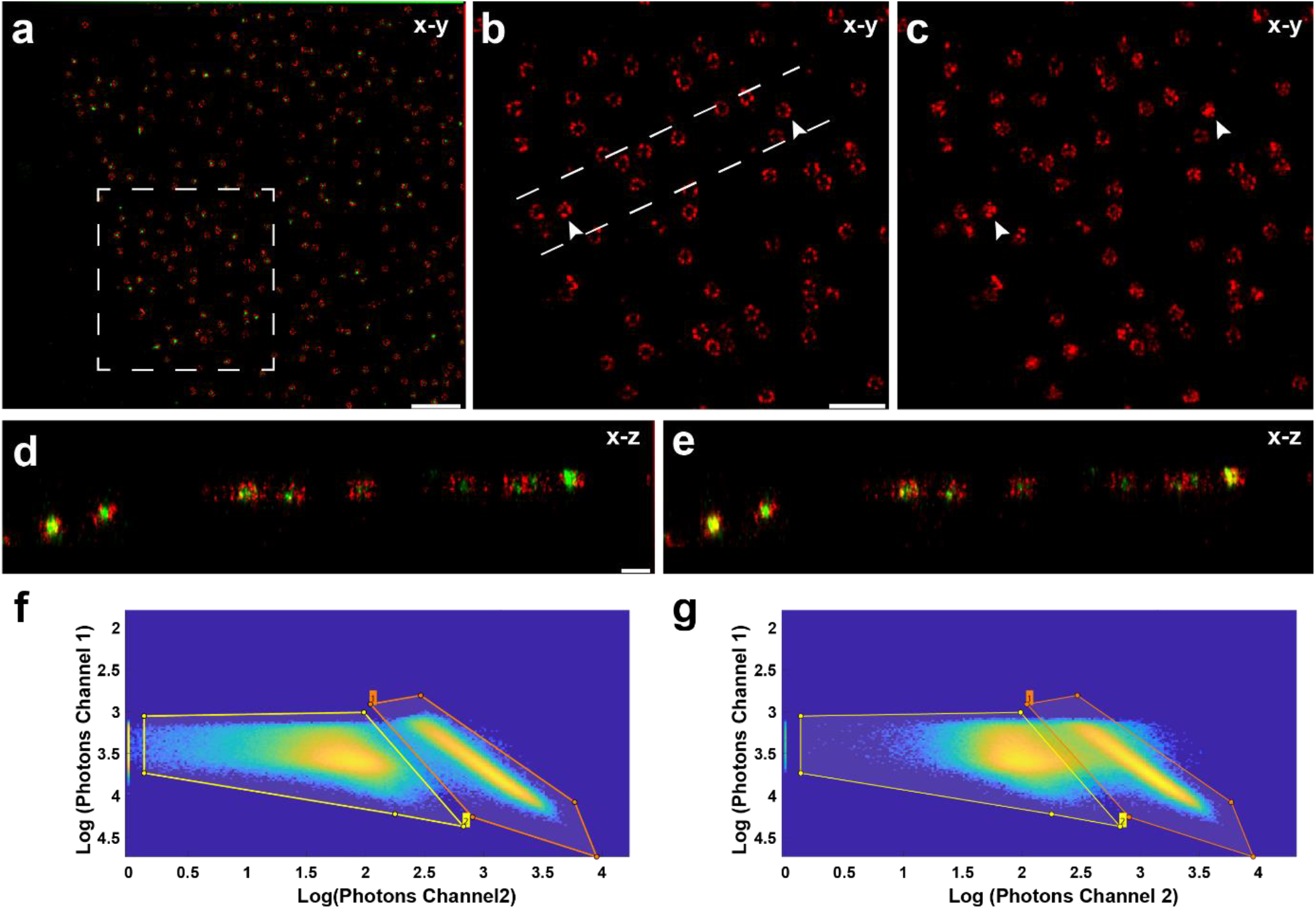
Ratiometric dual color 3D astigmatism imaging of Nup96-SNAP-AF647 and WGA-CF680. **a**, Dual color image of Nup96 (red) and WGA (green) in a U2OS cell. Nup96 image of the boxed region in **a** reconstructed by global fit (**b**) and individual fit (**c**) separately. Arrows indicate cross-talk of the reconstructed Nup96 image contaminated by WGA. **d** and **e** are the side-view of the region bounded by dashed lines in **b** reconstructed by global fit and individual fit separately. **f** and **g** are scatter plots of the photons in bright channel versus dark channel analyzed by global fit and individual fit separately. The boxed polygons are used to assign single molecules in different colors. Scale bars: 1 μm (**a**), 500 nm (**b**), 100 nm (**d**).

**Supplementary Fig. 9.**
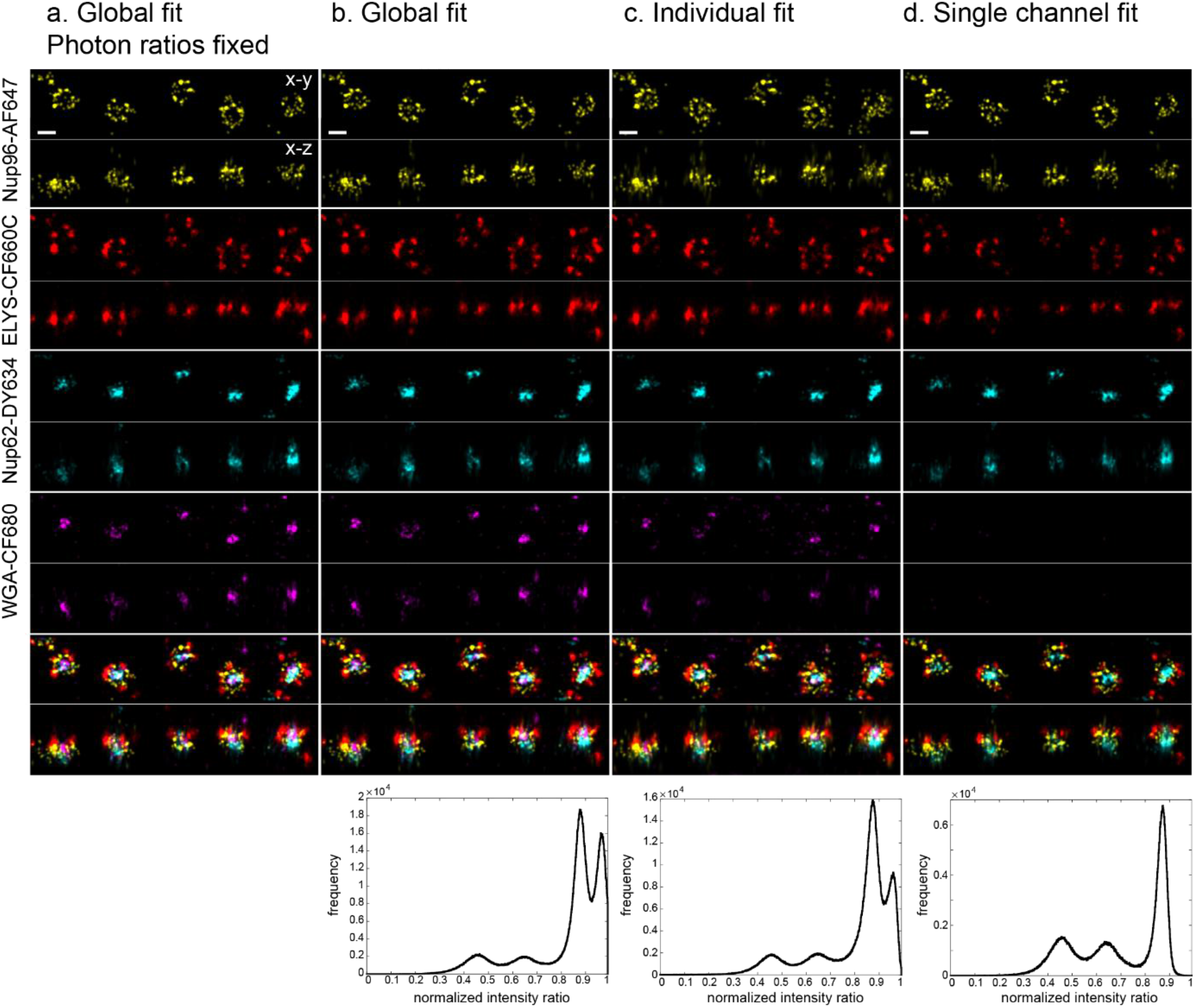
4 color 3D astigmatism imaging of Nup62-DY634, Nup96-AF647, ELYS-CF660C and WGA-CF680 using different reconstruction algorithms. **a**, Single molecule images are fitted with global fit with fixed photon ratios and the color was assigned based on the maximum likelihood. **b**, Global fit with only linking xyz and extracting the color from the ratio of fitted intensities in both channels. **c**, Individual fit of each channel and color assignment as in **b**. Here, candidate peaks from either channel are fitted also in the other channel. **d**, Individual fit of each channel, followed by linking corresponding localizations and color assignment as in b. As CF680 is hardly ever detected in Channel 2, it disappears from the image. Scale bars 100 nm.

**Supplementary Fig. 10.**
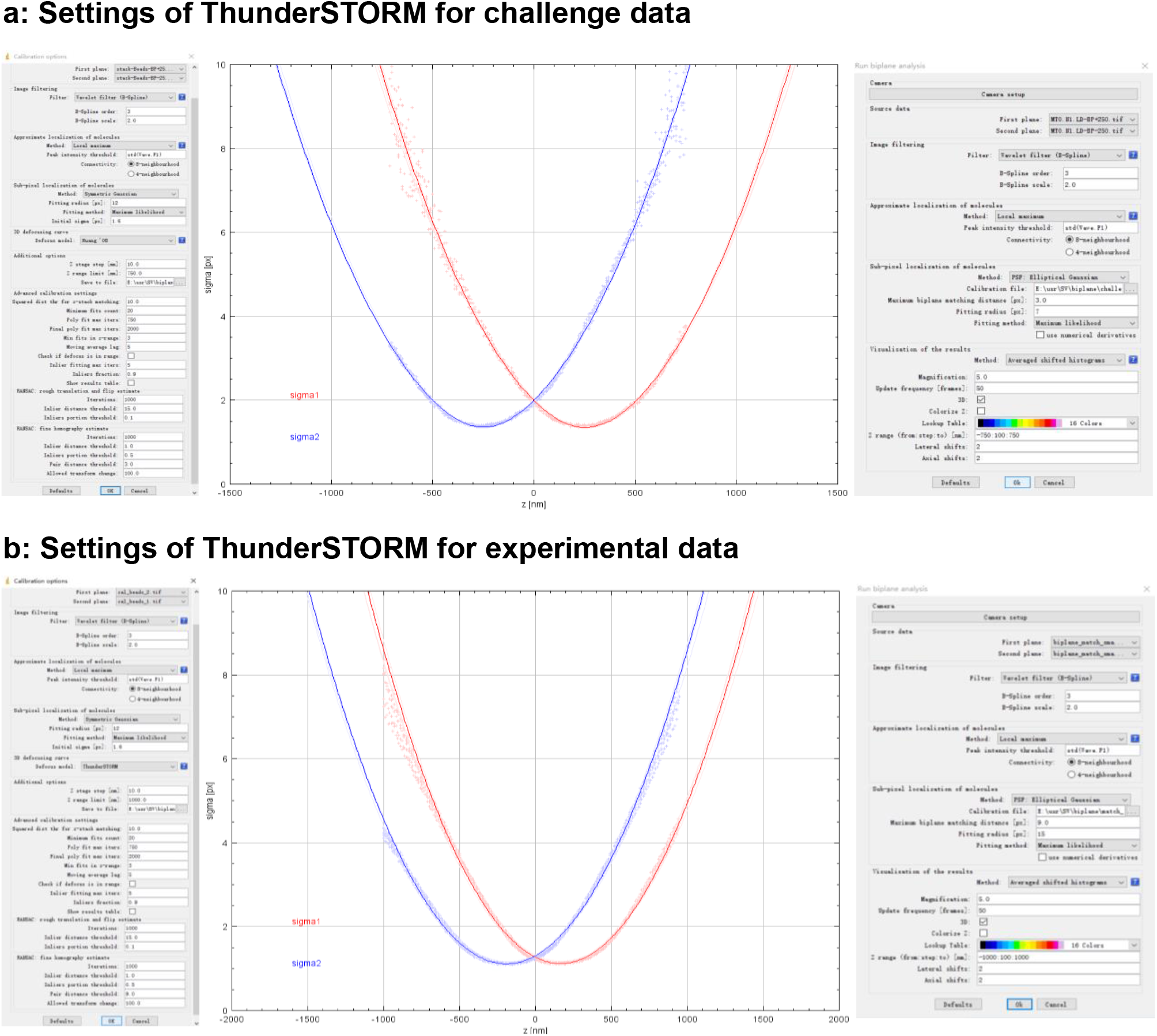
Settings of ThunderSTORM software used for calibrating and analyzing biplane data. **a**. Left to right panel: Calibration settings of the 2016 SMLM Challenge 3D-Biplane data ranging from -750 nm to 750 nm in the axial direction, the corresponding calibration curve and analysis settings of the challenge training biplane datasets (MT0. N1. LD_BP). The reconstructed SMLM image is shown in **Figure 1e4**. **b**. Left to right panel: Calibration settings of an experimental *z*-stack beads ranging from -1 μm to 1 μm in the axial direction, the corresponding calibration curve and analysis settings of the experimental biplane datasets (Nup96-AF647). The reconstructed SMLM image is shown in **Figure 2a4**.

**Supplementary Table 1.**

|  | **parameter** | **challenge data** | **experimental data** |
| --- | --- | --- | --- |
| Calibration parameters | Fitting radius[px] | 12 | 12 |
|  | Defocus model | Huang'08 | ThunderSTORM |
|  | Z range limit[nm] | 750 | 1000 |
|  | Pair distance threshold | 3 | 9 |
| Biplane analysis parameters | Maximum biplane matching distance[px] | 3 | 9 |
|  | Fit radius[px] | 7 | 15 |
|  | Z rang(from:step:to)[nm](3D) | -750:100:750 | -1000:100:1000 |

Calibration and analysis parameters that are different from the default values of ThunderSTORM software (ThunderSTORM version: dev-2016-09-10-b1, github.com/zitmen/thunderstorm/releases/tag/dev-2016-09-10-b1).

### Supplementary Note 1

#### Parameter Merging for individual fits of multi-channels

To merge parameters returned from individual fits of different channels, weighted arithmetic mean of parameters from all channels was used. Here, we used the reciprocal of the estimated CRLB as the weights for each parameter:

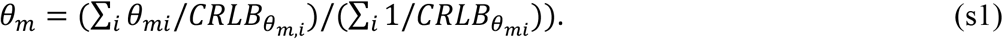

Here, *θ*_*mi*_ is the set of parameters being estimated in the *i*_th_ channel and 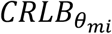 is the corresponding CRLB. The significance of this choice is that this weighted mean is the maximum likelihood estimator of parameters of different channels under the assumption that they are independent and normally distributed with the same mean. Therefore, this combination could return the optimized localization precision. As show in **Supplementary Fig. 1a** and **Supplementary Fig. 2a**, the localization accuracy of the global fit with linking *xyz* is similar to that of the CRLB-weighted individual fits. However, the global fit could substantially improve the localization accuracy in *z* by additionally linking photon parameters.

### Supplementary Note 2

#### Calculation of multichannel CRLB

To quantify the localization precision of globLoc fitter, we compared it with the CRLB which is the limiting lower bound of the variance for any unbiased estimator. The general definition of CRLB is evaluated as the diagonal element of the inverse of the Fisher information matrix:

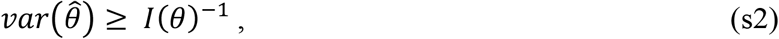

where 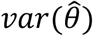 is the variance of an estimator and *I*(*θ*) is the Fisher information matrix. Depending on how the parameters are linked during fit, the Fisher information matrix is defined as

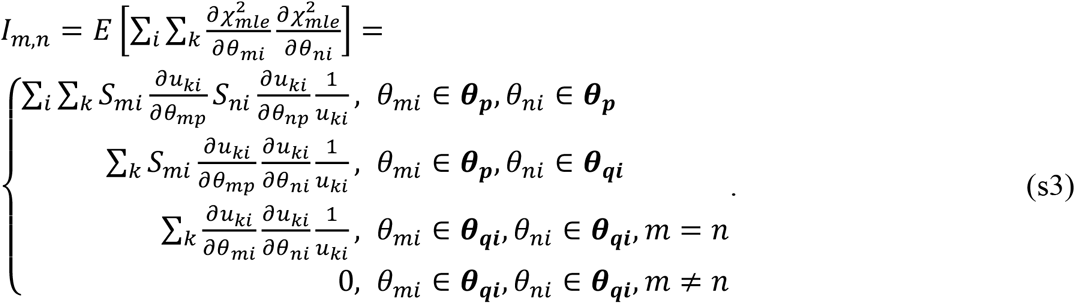

Here, 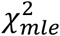 is defined as Equation (1) in Methods, *μ*_*ki*_ is the expected photon number in the *k*th pixel of the *i*th channel. ***θ***_***p***_ is the set of global parameter and ***θ***_***qi***_ is the set of local parameters of *i*th channel.

### Supplementary Note 3

#### Derivatives for IAB-based 4Pi-PSF model

The IAB-based 4Pi-PSF model is written as in ref. ^25^:

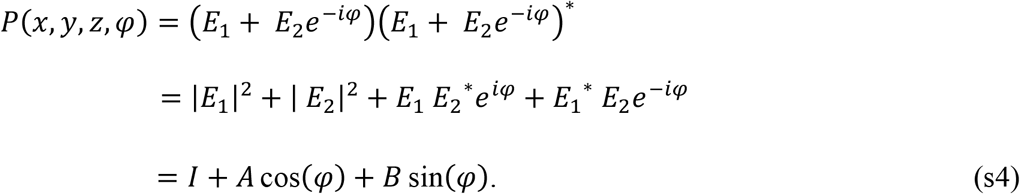

Here, *φ* is the interference phase. *I*(*x, y, z*), ***A***(*x, y, z*) and ***B***(*x, y, z*) are phase independent and slowly varying real functions of *x, y, z*. In order to construct the Hessian and Jacobian matrix, the following partial derivatives of *x, y, z*, phase, photons and background are used:

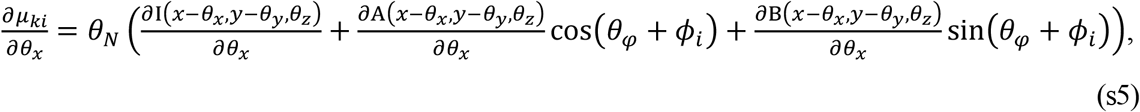

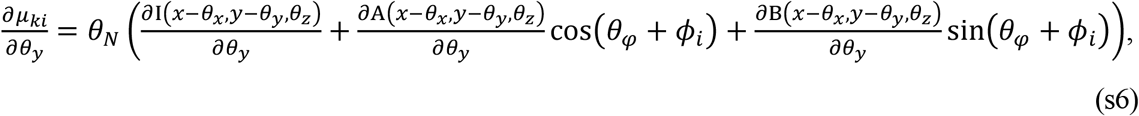

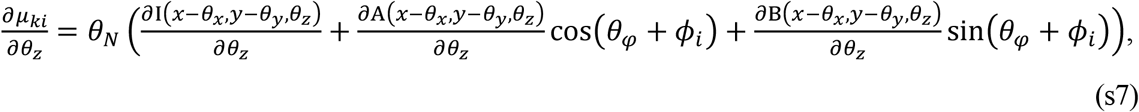

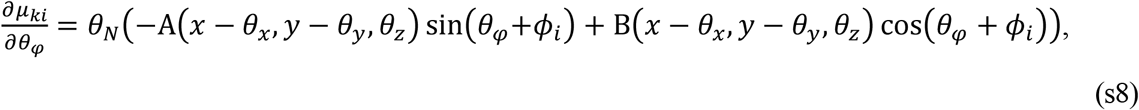

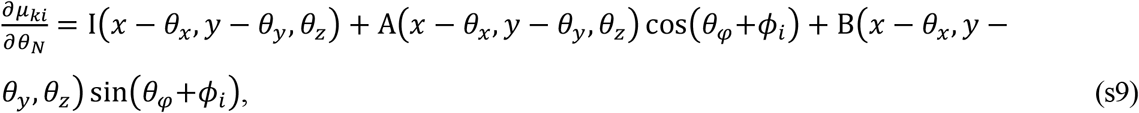

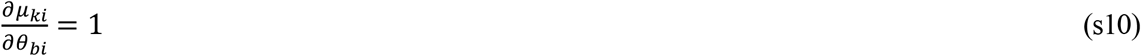

Here, the cubic splines are used to interpolate the 3D matrices *I, A* and *B* to calculate the partial derivative along *x, y* and *z*, separately^21,30^. The L-M iterative process is considered to be converged when the ratio of the relative change of 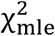 is less than 10^−6^ compared to the last iteration.

### Supplementary Note 4

#### Method for L-M nonlinear optimization of multichannel data

For maximum likelihood estimation, the cost function 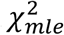 is defined as Equation (1) in Methods. During the L-M optimization process, Hessian (*H*_*m,n*_) and Jacobian (*J*_*m*_) matrix are defined as Equation (4) and (5) in the Methods. The detailed optimization algorithm used in this work is described below:

1. Calculate 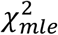 (old) with a user defined starting parameters *θ*_*m*_.
2. Initialize λ as 0.1. λ is a damping factor that controls whether the L-M fit should behave more as a gradient descent fit method (lambda<<1) or an expansion fit method (lambda>>1).
3. Calculate the updates of each parameter *Δθ*_*m*_ by solving the linear equations:

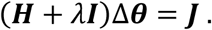
4. Derive new trial-fit-parameters *θ*_*m*_(*new*) as follows:

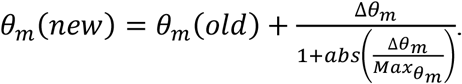

Here, 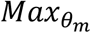 is a clamping factor which controls the maximum change of parameter *θ*_*m*_ during each iteration. If the sign of *Δθ*_*m*_ has changed since last update, 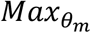 is multiplied by 0.5 to suppress oscillations during optimization and damp excessively large corrections^33^.
5. Determine 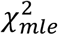 (new) with the new parameter using Equation (1) in the Methods.
6. If 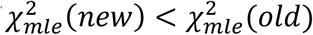, substitute λ with λ/10, and *θ*_*m*_(*old*) with *θ*_*m*_(*new*) and continue with step 3.
7. If 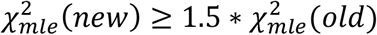, substitute λ with 10λ, keep *θ*_*m*_(*old*) unchanged, and continue with step 3.
8. If 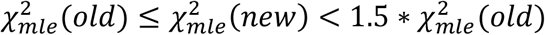, keep λ unchanged, substitute *θ*_*m*_(*old*) with *θ*_*m*_(*new*), and continue with step 3.

This iterative calculation may be stopped when one of the following conditions is met:

a. The ratio of the relative change of 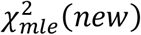 and 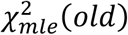 decreases to less than a specified value (10^−6^ in this work).

The iteration time of the calculation loops exceeds a maximum number.

## REFERENCES

1. Bossi, M. et al. Multicolor far-field fluorescence nanoscopy through isolated detection of distinct molecular species. Nano Lett. 8, 2463–2468 (2008).

2. Mund, M. et al. Systematic Nanoscale Analysis of Endocytosis Links Efficient Vesicle Formation to Patterned Actin Nucleation. Cell 174, 884–896 (2018).

3. Zhang, Y. et al. Nanoscale subcellular architecture revealed by multicolor threedimensional salvaged fluorescence imaging. Nat. Methods 17, 225–231 (2020).

4. Zhang, Z., Kenny, S. J., Hauser, M., Li, W. & Xu, K. Ultrahigh-throughput singlemolecule spectroscopy and spectrally resolved super-resolution microscopy. Nat. Methods 12, 935–938 (2015).

5. Juette, M. F. et al. Three-dimensional sub–100 nm resolution fluorescence microscopy of thick samples. Nat. Methods 5, 527–529 (2008).

6. Babcock, H. P. Multiplane and spectrally-resolved single molecule localization microscopy with industrial grade CMOS cameras. Sci. Rep. 8, 4–11 (2018).

7. Jia, S., Vaughan, J. C. & Zhuang, X. Isotropic three-dimensional super-resolution imaging with a self-bending point spread function. Nat. Photonics 8, 302–306 (2014).

8. Dasgupta, A. et al. Direct supercritical angle localization microscopy for nanometer 3D superresolution. Nat. Commun. 12, 1180 (2021).

9. Cabriel, C. et al. Combining 3D single molecule localization strategies for reproducible bioimaging. Nat. Commun. 10, 1–10 (2019).

10. Shtengel, G. et al. Interferometric fluorescent super-resolution microscopy resolves 3D cellular ultrastructure. Proc. Natl. Acad. Sci. 106, 3125–3130 (2009).

11. Huang, F. et al. Ultra-High Resolution 3D Imaging of Whole Cells. Cell 166, 1028–1040 (2016).

12. Chen, L. et al. Advances of super-resolution fluorescence polarization microscopy and its applications in life sciences. Comput. Struct. Biotechnol. J. 18, 2209–2216 (2020).

13. Gu, L. et al. Molecular resolution imaging by repetitive optical selective exposure. Nat. Methods 16, 1114–1118 (2019).

14. Cnossen, J. et al. Localization microscopy at doubled precision with patterned illumination. Nat. Methods 17, 59–63 (2020).

15. Jouchet, P. et al. Nanometric axial localization of single fluorescent molecules with modulated excitation. Nat. Photonics 15, 297–304 (2021).

16. Gu, L. et al. Molecular-scale axial localization by repetitive optical selective exposure. Nat. Methods 18, 369–373 (2021).

17. Ram, S., Prabhat, P., Chao, J., Ward, E. S. & Ober, R. J. High accuracy 3D quantum dot tracking with multifocal plane microscopy for the study of fast intracellular dynamics in live cells. Biophys. J. 95, 6025–6043 (2008).

18. Liu, S., Kromann, E. B., Krueger, W. D., Bewersdorf, J. & Lidke, K. A. Three dimensional single molecule localization using a phase retrieved pupil function. Opt. Express 21, 29462–29487 (2013).

19. Wolter, S. et al. rapidSTORM: accurate, fast open-source software for localization microscopy. Nat. Methods 9, 1040–1041 (2012).

20. Baddeley, D. et al. 4D super-resolution microscopy with conventional fluorophores and single wavelength excitation in optically thick cells and tissues. PLoS One 6, e20645(2011).

21. Li, Y. et al. Real-time 3D single-molecule localization using experimental point spread functions. Nat. Methods 15, 367–369 (2018).

22. Sage, D. et al. Super-resolution fight club: assessment of 2D and 3D single-molecule localization microscopy software. Nat. Methods 16, 387–395 (2019).

23. Ovesný, M., Křížek, P., Borkovec, J., Švindrych, Z. & Hagen, G. M. ThunderSTORM: a comprehensive ImageJ plug-in for PALM and STORM data analysis and superresolution imaging. Bioinformatics 30, 2389–2390 (2014).

24. Thevathasan, J. V. et al. Nuclear pores as versatile reference standards for quantitative superresolution microscopy. Nat. Methods 16, 1045–1053 (2019).

25. Li, Y. et al. Accurate 4Pi single-molecule localization using an experimental PSF model. Opt. Lett. 45, 3765 (2020).

26. Lehmann, M., Lichtner, G., Klenz, H. & Schmoranzer, J. Novel organic dyes for multicolor localization-based super-resolution microscopy. J. Biophotonics 9, 161–170 (2016).

27. Wu, Y. et al. Maximum-likelihood model fitting for quantitative analysis of SMLM data. 1–39 (2021) doi:10.1101/2021.08.30.456756.

28. Ries, J. SMAP: a modular super-resolution microscopy analysis platform for SMLM data. Nat. Methods 17, 870–872 (2020).

29. Laurence, T. A. & Chromy, B. A. Efficient maximum likelihood estimator fitting of histograms. Nat. Methods 7, 338–339 (2010).

30. Babcock, H. P. & Zhuang, X. Analyzing Single Molecule Localization Microscopy Data Using Cubic Splines. Sci. Rep. 7, 552 (2017).

31. Stallinga, S. & Rieger, B. Accuracy of the Gaussian Point Spread Function model in 2D localization microscopy. Opt. Express 18, 24461 (2010).

32. Deschamps, J., Rowald, A. & Ries, J. Efficient homogeneous illumination and optical sectioning for quantitative single-molecule localization microscopy. Opt. Express 24, 28080–28090 (2016).

33. Tobergte, D. R. & Curtis, S. Daophot: a Computer Program for Crowded-Field Stellar Photometry (Psf Photometry). J. Chem. Inf. Model. 53, 1689–1699 (2013).

