## Supplementary material for "Global fitting for high-accuracy multi-channel single-molecule localization": Tutorial of globFit

### GlobFit examples: tutorial

### Getting started

Here, we show you how to use global fitting in SMAP. You find a link to the Matlab and compiled versions to SMAP as well as the documentation and example data at: [www.rieslab.de/#software](www.rieslab.de/#software).

Please follow the SMAP *User Guide* for installation and detailed description of general workflows. We also recommend you to follow the *Getting Started Guide* with examples

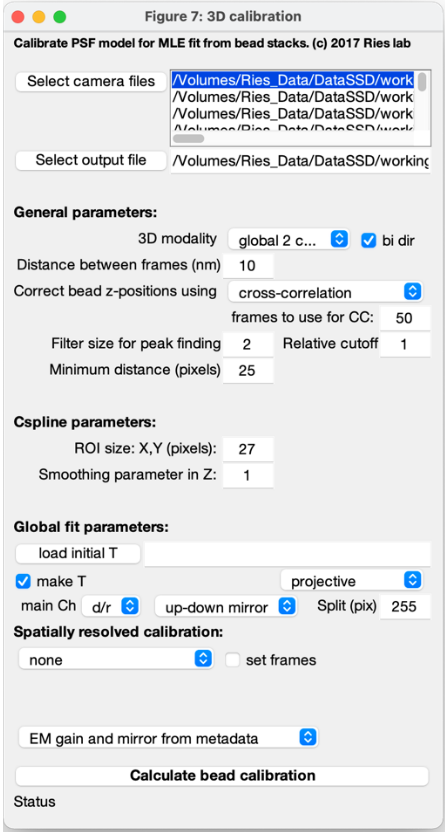

*Figure 1: Settings for the plugin: calibrate3DsplinePSF.*

### Example: 4 color 3D Nuclear pore complex

The example files for globFit you can find at: [https://www.embl.de/download/ries/globLoc/](https://www.embl.de/download/ries/globLoc/) .To analyze the 4-color 3D nuclear pore complex (NPC) data, please download and extract the NPC4C.zip. Please note that this is only a subset of the frames used to for Figure 2 in the manuscript.

#### Bead calibration

Compare section 5.4 in the *User Guide*.

1. Start SMAP
2. Select the **calibrate3DsplinePSF** plugin from the **Menu/Analyze/sr3d** or in the **Analyze/sr3D** tab and press Run.
3. Load bead images with select camera files:
    a. You can add individual bead stacks individually with add or add an entire directory containing many bead stacks with add dir.
    b. Set the following parameters according to Figure 1.
    c. Calculate bead calibration.

#### Global fitting with SMAP

For general settings (single-channel fitting) that are also relevant for globFit see the *User Guide* section 5.

4. In the localize tab in the bottom press Change to load the workflow: `fit_global_dualchannel.txt`
5. load images: select any of the individual image tiff files.
6. Check if metadata has been set by clicking on set Cam Parameters. Importantly, the coordinates of the ROI need to be: [100, 0, 287, 512].
7. In the **Peak Finder** tab load the transformation with load T and select the bead calibration file that you created before, i.e. `beads_3dcal.mat`.
8. In the **Fitter** tab load 3D cal and select the same file again.
9. Set the parameters according to Figure 2.
10. You can select in the table on the right had side which parameters to link during the fit (here: x,y,z).
11. Check if fitting works with Preview. You can adjust the cutoff (**dynamic factor**) in the peak finder to fit more or less candidates.
12. Press Localize to start the fitting. Depending on the computer and if the GPU or CPU are used, this can take between a few minutes and an hour.

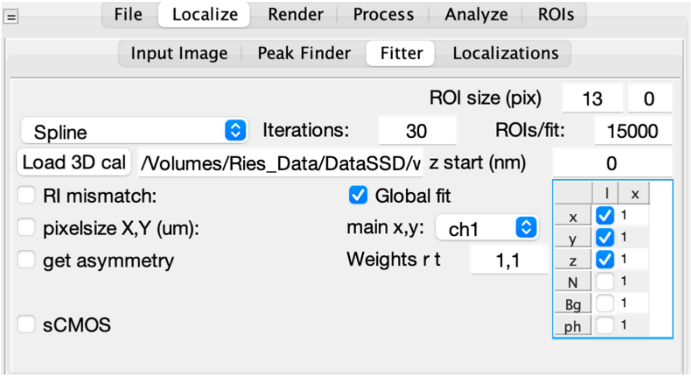

*Figure 2: Parameters for the fitting workflow.*

#### Post-processing and Color assignment

##### Rendering

When the fitting is done you can render the super-resolution image. To this end, press Render in the **Render** tab. Please consult the *User Guide* section 7 for detailed instructions and familiarize yourself with the rendering.

##### Filtering

Use the versatile filtering options of SMAP (*User Guide* section 6.2) to only display localizations of sufficient quality (we recommend filtering by the lateral localization precision to only display bright localization events, and by the relative log-likelihood (LLrel) to reject erroneous localizations from fluorophores activated too close to each other.

##### Drift correction

You can perform a 3D drift correction based on redundant cross-correlation as explained in the *User Guide* section 6.3.

##### Color assignment

Here we discuss, how to assign to each localization its color (compare *User Guide* section 9.2.2):
Open the plugin **Process/Assign2C/Intensity2ManyChannels.** This plugin assigns color values to the localizations based on two fields of the localization data, here the photon numbers fitted in channel 1 (phot1) and channel 2 (phot2), respectively. These values are assigned to the channel field of the localizations.

13. Select the two fields that encode the intensity in both channels, here phot1 and phot2. From this selection an image is generated in which the x-axis corresponds to the first intensity value and the y-axis corresponds to the second intensity value (logarithmic scaling). By checking log scale you can also use a logarithmic scaling for the contrast.

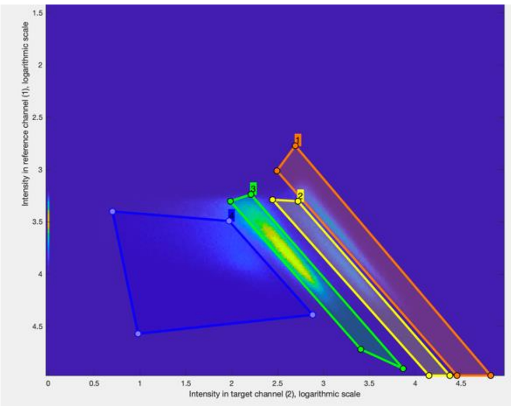

*Figure 3: Dual color assignment. ROIs used to assign four colors to the intensity ratios.*

14. Press ROI 1 to draw a polygon ROI around the area the image that encloses all localizations corresponding to Color 1 (Figure 3).
15. Repeat the same for Color 2 and additional Colors (for Color 4 and onwards put the color number in the respective field).
16. With Show ROIs you can display all ROIs. You can always adjust all ROIs.
17. With Delete ROIs you can delete all ROIs and start over.
18. With load and save you can load and save all defined ROIs for later use.
19. Press Run to assign a Color number to all localizations based on the defined ROIs.
    a. If use grouped is selected, this assignment is based on grouped localizations.
    b. Localizations outside of any ROI are assigned the channel value 0. This is also the channel value for all data before channel assignment.
20. In the **Render** tab please create 3 additional layers with the + tab.
    a. In Layer 1, put in 1 for the Color in the Ch field. For Layers 2-4 put the color values 2-4.
    b. In each layer, choose a different lookup table (LUT).
21. Now you should be able to render a four-color image.
22. In the 'layers' panel on the top-right side of the GUI you can select which layers to display, and also if to show the layers separately (split) and if to show in addition the composite image (comp).

##### 3D visualization

For 3D visualization with the SMAP 3D viewer please follow the *User Guide* section 7.6.

### Example: bi-plane SMLM of the nuclear pore complex

To analyze the bi-plane 3D nuclear pore complex (NPC) data, please download and extract the NPC_BP.zip from https://www.embl.de/download/ries/globLoc/.

#### Bead calibration

1. Start SMAP
2. Select the ***calibrate3DsplinePSF*** plugin from the ***Menu/Analyze/sr3d*** or in the ***Analyze/sr3D*** tab and press Run.

3. Select camera files to load all the bead stacks. A new window opens. Here press add dir and select all directories in the bead directory. Press Done
4. Set the parameters according to Figure 4.
    a. As we have bi-plane data in two parts of the camera we choose the 3D modality: global 2 channel.
    b. For nicer visualization of the validation of the model you can select • bi dir to activate bi-directional fitting.
    c. Set the distance between slices in the stack (in nanometers).
    d. You can adjust the other parameters according to the *User guide* section 5.4.1.
    e. An initial transformation is calculated from the bead stacks. You can choose the main channel (in this case the bottom channel) and specify that the upper and lower channels are mirrored horizontally with respect to each other.
    f. We let SMAP determine the EM gain and mirror from the metadata. Otherwise, set here if beads were acquired with the conventional or EM mode.
5. Calculate bead calibration. This step might take 5-10 minutes, you can see the progress in the status bar of the plugin. You can compare your saved files with beads_BP_3dcal.fig. Specifically, compare the tabs of the output figure with those in Figure 5.

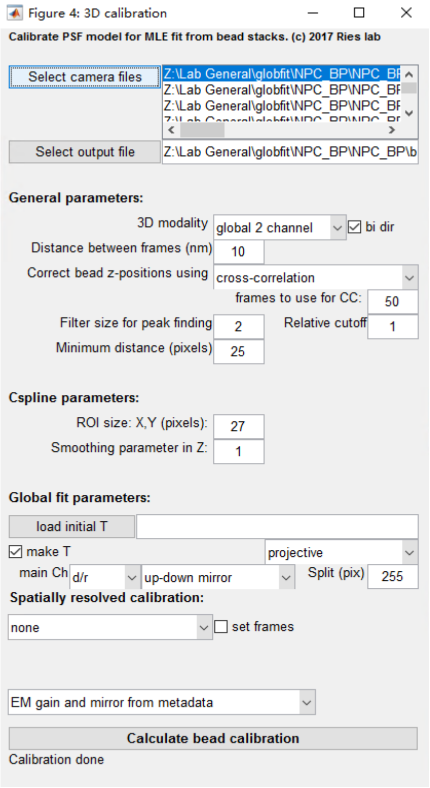

*Figure 4: Settings for the plugin: calibrate3DsplinePSF.*

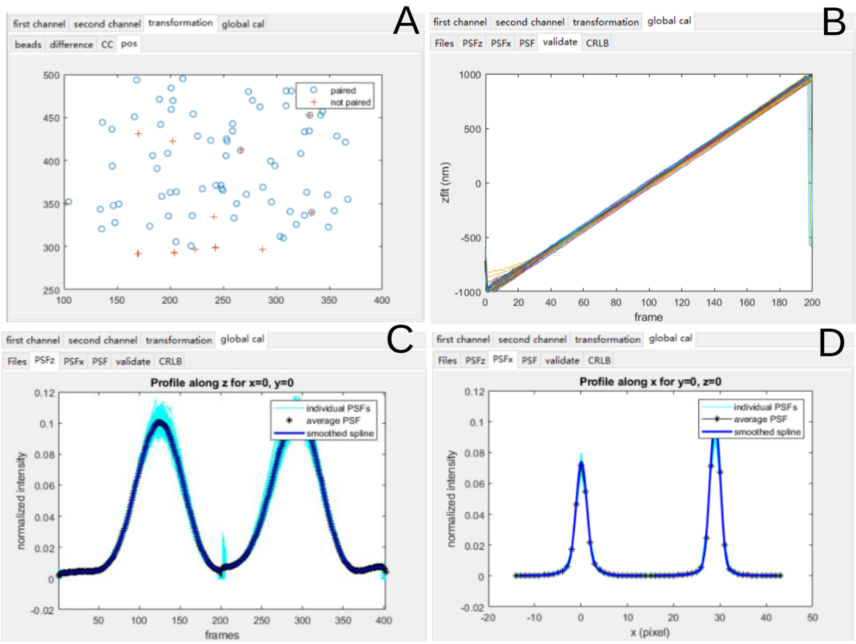

*Figure 5: Output of the GUI calibration. A) to check if beads across the whole field of view were paired. B) to check if the model could properly fit the individual beads. C) Cross-section along z-axis through model to check if the model represents the individual beads. D) as C. but along x-axis.*

#### Fit bi-plane SMLM data with SMAP

For general settings (single-channel fitting) that are also relevant for globFit see the User Guide section 5.

6. Check in the SMAP GUI at the bottom of the Localize tab that the fitting workflow fit_global_dualchannel is selected. Otherwise load that workflow by pressing Change.

7. In the main SMAP GUI select the **Localize/Input Image** tab.
8. Load images and select one of the files in the bi_plane data directory .The loader might display an **error message** that it cannot load the images. Ignore this for the moment and try the next steps. This error message comes from the micro-manager code, thus it cannot easily be fixed in SMAP.
9. Check if metadata has been set by clicking on set Cam Parameters. Importantly, the coordinates of the ROI need to be: [2, 0, 512, 512].
10. In the **Peak Finder** tab load the transformation with load T and select the bead calibration file that you created before, i.e. beads_BP_3dcal.mat.
11. In the **Fitter** tab load 3D cal and select the same file again.
12. You can select in the table on the right had side (figure 6) which parameters to link during the fit (here: x,y,z,N,Bg).
13. You can adjust the parameters in the **Peak Finder** tab according to the *User guide* section 5.3.
14. Check if fitting works with Preview. You can adjust the cutoff (dynamic factor) in the peak finder to fit more or less candidates.
15. Press Localize to start the fitting. Depending on the computer and if the GPU or CPU are used, this can take between a few minutes and an hour.

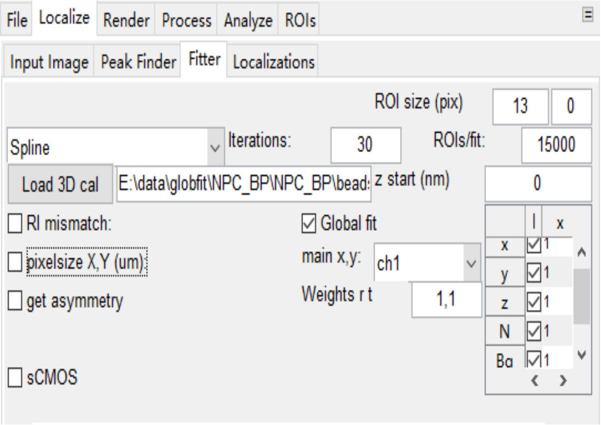

*Figure 6: Parameters for the fitting workflow*

#### Post-processing

##### Filtering

Use the versatile filtering options of SMAP (User Guide section 6.2) to only display localizations of sufficient quality (we recommend filtering by the lateral localization precision to only display bright localization events, and by the relative log-likelihood (LLrel) to reject erroneous localizations from fluorophores activated too close to each other).

##### Drift correction

You can perform a 3D drift correction based on redundant cross-correlation as explained in the User Guide section 6.3.

##### Rendering

You can render the super-resolution image in the Render tab. Please consult the User Guide section 7 for detailed instructions and familiarize yourself with the rendering.

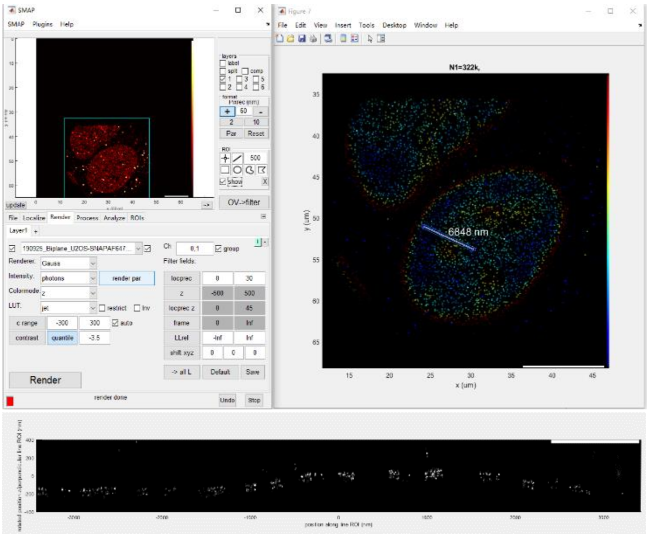

*Figure 7: GUI panels to render render the super-resolution image and side-view rendering of the localizations in the line ROI.*
